# Surface lipids influence developmental and species-specific chemical signalling in nematodes

**DOI:** 10.1101/2024.04.24.590549

**Authors:** Anna M. Kotowska, Fumie Hiramatsu, Morgan R. Alexander, David J. Scurr, James W. Lightfoot, Veeren M. Chauhan

**Affiliations:** Advanced Materials and Healthcare Technologies Division, School of Pharmacy, University of Nottingham, University Park Nottingham, NG7 2RD, UK; Max Planck Research Group Genetics of Behavior, Max Planck Institute for Neurobiology of Behavior – caesar, Ludwig-Erhard-Allee 2, 53175, Bonn, Germany

**Keywords:** Nematodes, Lipids, 3D-OrbiSIMS, Signalling, β-oxidation, *daf-22*, Metabolism, Predation, Development, Evolution

## Abstract

Chemical signalling facilitates organismal communication and coordinates physiological and behavioural processes. In nematodes, signalling has predominantly focused on secreted molecules leaving the surface’s communicative potential unexplored. By utilising 3D-OrbiSIMS, an advanced surface-sensitive mass spectrometry method, we directly characterised the molecular composition of the outermost regions (50 nm in depth) of *Caenorhabditis elegans* and *Pristionchus pacificus* to improve the understanding of surface-mediated chemical communication. We found that nematode surfaces consist of a lipid-dominated landscape (> 81 % *C. elegans* and > 69 % *P. pacificus* of all surveyed chemistries) with distinct compositions, which enrich in granularity and complexity through development. The surface-anchored lipids are also species-specific, reflecting evolutionary and ecological adaptations to their environmental niches. By exploring the effect of mutations on lipid production we found the peroxisomal fatty acid β-oxidation component *daf-22* is essential for defining the surface molecular fingerprint. This pathway is conserved across species in producing distinct chemical profiles, indicating its fundamental role in lipid metabolism and for maintaining surface integrity and function. Furthermore, we discovered that variations in surface-anchored lipids of *C. elegans daf-22* larvae contribute to significantly increased susceptibility to predation by *P. pacificus*. Therefore, our findings reveal that the nematode surface is not just a passive boundary but a dynamic signalling platform with evolved, species-specific signatures. These molecular mechanisms are pivotal in shaping identity and communication strategies, providing new insights into the evolutionary and ecological dynamics of chemical signalling across organisms.

**HIGHLIGHTS:**

- Nematode surfaces are lipid-rich, changing with development.
- Surface lipids are species-specific, reflecting adaptations.
- The *daf-22* gene is key in shaping the surface lipid profile.
- Surface lipids mediate predator-prey interactions in nematodes.

## INTRODUCTION

Organisms produce a multitude of chemical signals that can be secreted into their surroundings or displayed on their surface ^1^. These signals relay important physiological and environmental information to con-specifics and other co-habitants. However, due to the blend of different chemistries, deciphering the contribution of specific components can be challenging. The nematode *Caenorhabditis elegans*, with its advanced genetic and molecular tools, serves as an important model for understanding chemical signalling^2^. This has enabled the biological activity and molecular mechanisms involved in the synthesis of a variety of chemical cues to be elucidated. For example, a suite of volatile small molecule pheromones, known as ascarosides ^3^, have been found to be secreted into the nematode surroundings and regulate diverse biological processes ^4^. This encompasses regulating development, such as entering the stress-resistant dauer stage, and behaviours, including mate attraction, foraging and aggregation ^5^. However, it is likely that these secreted long-range signals are only one facet of the nematode’s communication strategy.

The nematode cuticle and its surface coat represent the outermost layer, connecting them with their external environment. In free-living nematodes, the cuticle acts as a permeability barrier ^6^ and must protect these organisms against both external abiotic hazards such as desiccation ^7^, as well as biological factors including pathogenic bacteria ^8^, fungal traps ^9^ and predatory nematodes ^10^. The cuticle comprises crosslinked collagen patterns ^11^, glycoproteins, cuticulins, and lipids ^12^, which are synthesised by the hypodermal cells ^13^. Furthermore, as the nematodes undergo larval moults, their surface coat is replenished and replaced. The expression of many of these components are tightly regulated and oscillate in synchronicity with the organism’s development and moult cycle ^14^. Therefore, this sophisticated surface architecture underpins not only the mechanical properties of these organisms, but also its chemical landscape, which plays a crucial role in behaviours, such as locomotion and mating interactions ^15^. Despite the identification of several surface proteins ^16^, the exact chemical composition and the broader significance of the nematode surface as a communication medium remain largely uncharted. This knowledge gap arises not from an absence of curiosity, but from the limitations in available tools for accurately capturing, analysing, and interpreting molecular surface chemistry.

## RESULTS

### Development influences surface lipids

The study of nematode surfaces has traditionally relied upon methods such as liquid chromatography-mass spectrometry (LC-MS) for analysing homogenates and surface extractions ^17^. Time-of-flight (TOF) based mass spectrometry techniques like Matrix-Assisted Laser Desorption/Ionization (MALDI) ^18, 19^ and Secondary Ion Mass Spectrometry (SIMS) ^19, 20^ have also been used to directly analyse surfaces, although these provide relatively lower mass accuracy. Advancements in surface-sensitive mass spectrometry, such as the 3D-OrbiSIMS, which combines a Gas Cluster Ion Beam (GCIB, Ar3000+) with an Orbitrap analyser ^21^, provide a significant uplift in the ability to understand the chemical complexity of biological samples ^22^, enabling direct surface chemical mapping with relatively high spatial resolution (≥ 2 µm) and mass resolving power (>240,000 at *m/z* 200), achieved in the absence of chemical fixation or additional labelling. Additionally, its field of view (500 µm × 500 µm), facilitates the imaging of the entire length of the nematode, enabling a comprehensive analysis of the organism’s chemical composition (**Fig. 1A**). Using this technique, the focus was to investigate the composition of the *C. elegans* surfaces, which consists of rich topological features that generate the organism’s external morphology (**Fig. S1**). Through control of the ion dose (3.00 × 10^14^ ions/cm^2^), the surface analysis was confined to the outermost regions (approximately 50 nm in depth ^23^), corresponding to the cortical cuticle of *C. elegans* ^24^. Global analysis of adult and newly hatched L1 larvae mass spectra suggested that their surfaces are very similar (**Fig. 1B, Fig. S2**). Distribution of molecular assignments into chemical classes (**Table S1**), revealed the OrbiSIMS spectra for both the adult and larvae *C. elegans* cuticle coats were dominated by surface-anchored lipids. Specifically, the total lipid composition, which includes fatty acids & triglycerides, phospholipids, ceramides, and sterols, contributed to 83.11% and 81.75% of the OrbiSIMS spectra for adults and larvae, respectively. Principal Component Analysis (PCA, **Fig. S3**) and a hierarchical clustering heatmaps (**Fig. 1D**) delineated the developmental stages of *C. elegans* and also initiated the categorisation of distinct chemical profiles responsible for the observed differences in surface chemistry.

**Figure 1.**
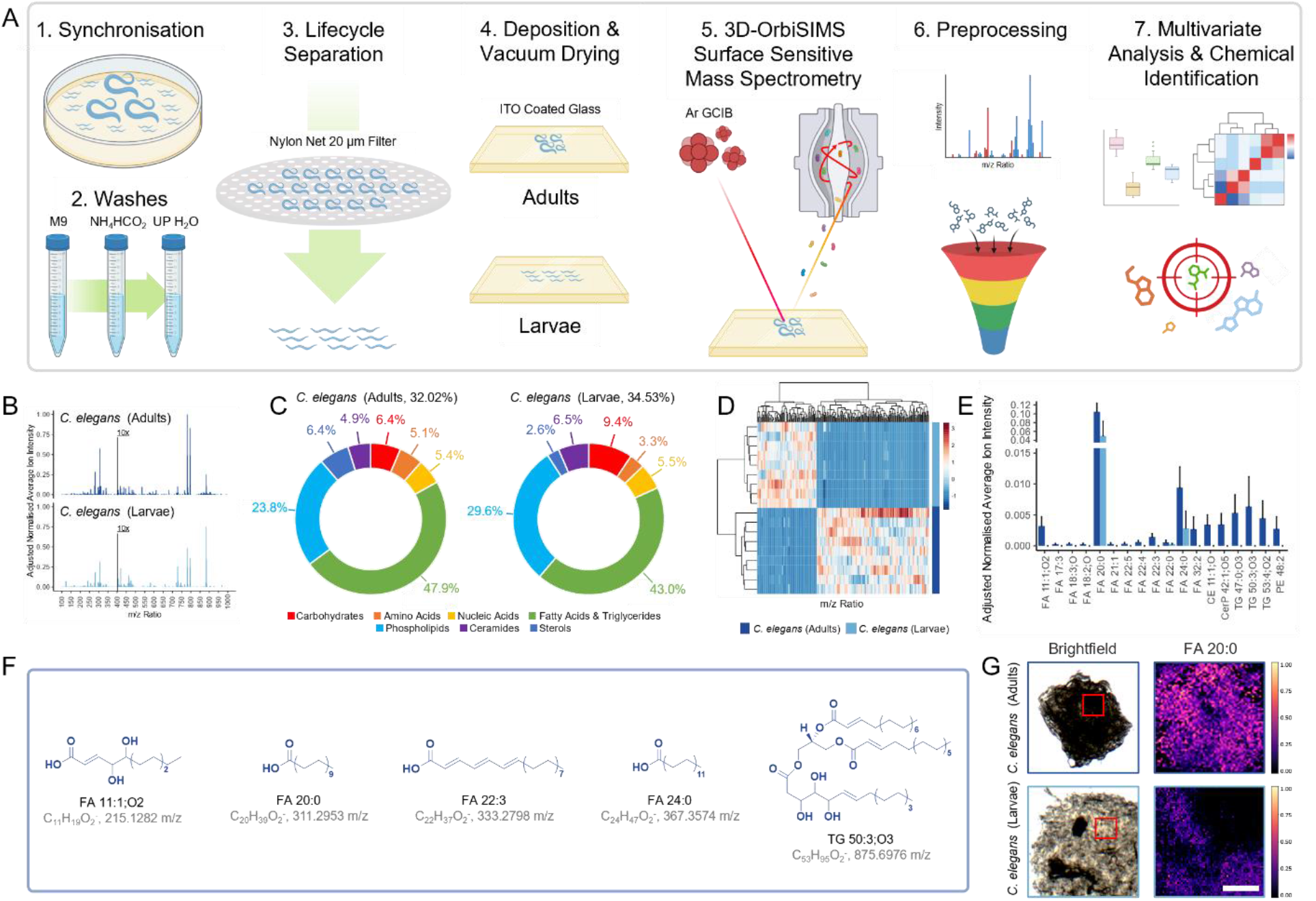
Surface specific chemistry for *C. elegans* adults and larvae evaluated using 3D-OrbiSIMS. **A** Schematic detailing capture of adults or larvae surface chemical maps using 3D-OrbiSIMS (100 µm^2^, n=9, created with BioRender.com). **B** Averaged *C. elegans* adults and larvae surface secondary ion mass spectra, normalised to maximum intensity across spectra, where intensity *m/z* >400 enhanced 10x for visibility. **C** Distribution of molecular assignments determined using chemical filtration (**Table S1**), as a percentage of total ions surveyed. **D** Hierarchical clustering heatmap for *C. elegans* adults and larvae *m/z* ratio intensity, for PC1 loadings with a standard deviation greater than the mean. **E** Significantly different chemistries on *C. elegans* adults and larvae surfaces (P<0.001 by Student’s t-test, n=9) present in LIPIDS MAPS (M-H, <2 ppm) with putative **F** chemical assignments h and structures. **G** Representative normalised intensity maps of *C. elegans* adult and larva surface chemistry, scale = 100 µm.

Secondary ions exhibiting significant intensity differences across developmental stages (*P*<0.001, Student’s t-test, n=9) were evaluated using the LIPID Metabolites And Pathways Strategy (LIPID MAPS) database ^25^. This facilitated the identification of molecular species and putative structural assignments (**Fig. 1E**) with a mass accuracy of < 2 ppm. Overall, there was a notable increase in the complexity of surface chemistries in adults, characterised by the prevalence of longer-chain fatty acids and triglycerides, which were seldom detectable in larvae. While certain fatty acids, such as FA 11:1;O2, 20:0, 22:0, and 24:0 (**Fig. 1F**), were identified in both developmental stages, their intensities remained significantly higher on the surface of adults. Representative normalised intensity maps visually highlight the differences in surface chemistry between *C. elegans* adults and larvae, with FA 20:0 displaying the most pronounced relative intensity across the two developmental stages (**Fig. 1G**). These observations, obtained using 3D-OrbiSIMS, provide important mechanistic insights into the dynamic and regulated nature of surface composition, highlighting the critical role of surface lipids in developmental transitions, which may also be important for relaying developmental stage-specific signals.

### Exploring evolutionary adaptations

While *C. elegans* is predominantly found in rotting fruit and is the most well-studied nematode species ^26^, the phylum Nematoda boasts a wide diversity, and encompasses species that have adapted to a variety of ecological niches. For instance, *Pristionchus pacificus*, is a nematode distantly related to *C. elegans* and is, frequently associates with scarab beetles ^27^. Importantly, the surface properties of nematodes, being in direct contact with their environment, are likely to reflect evolutionary adaptations essential for thriving in specific habitats. For example, *P. pacificus* displays a unique topological surface arrangement, when compared to *C. elegans* (**Fig. S1**). Therefore, we conducted direct surface analysis of *P. pacificus* using 3D-OrbiSIMS to explore potential species-specific chemical signatures (**Fig. 2A Fig. S4**).

**Figure 2.**
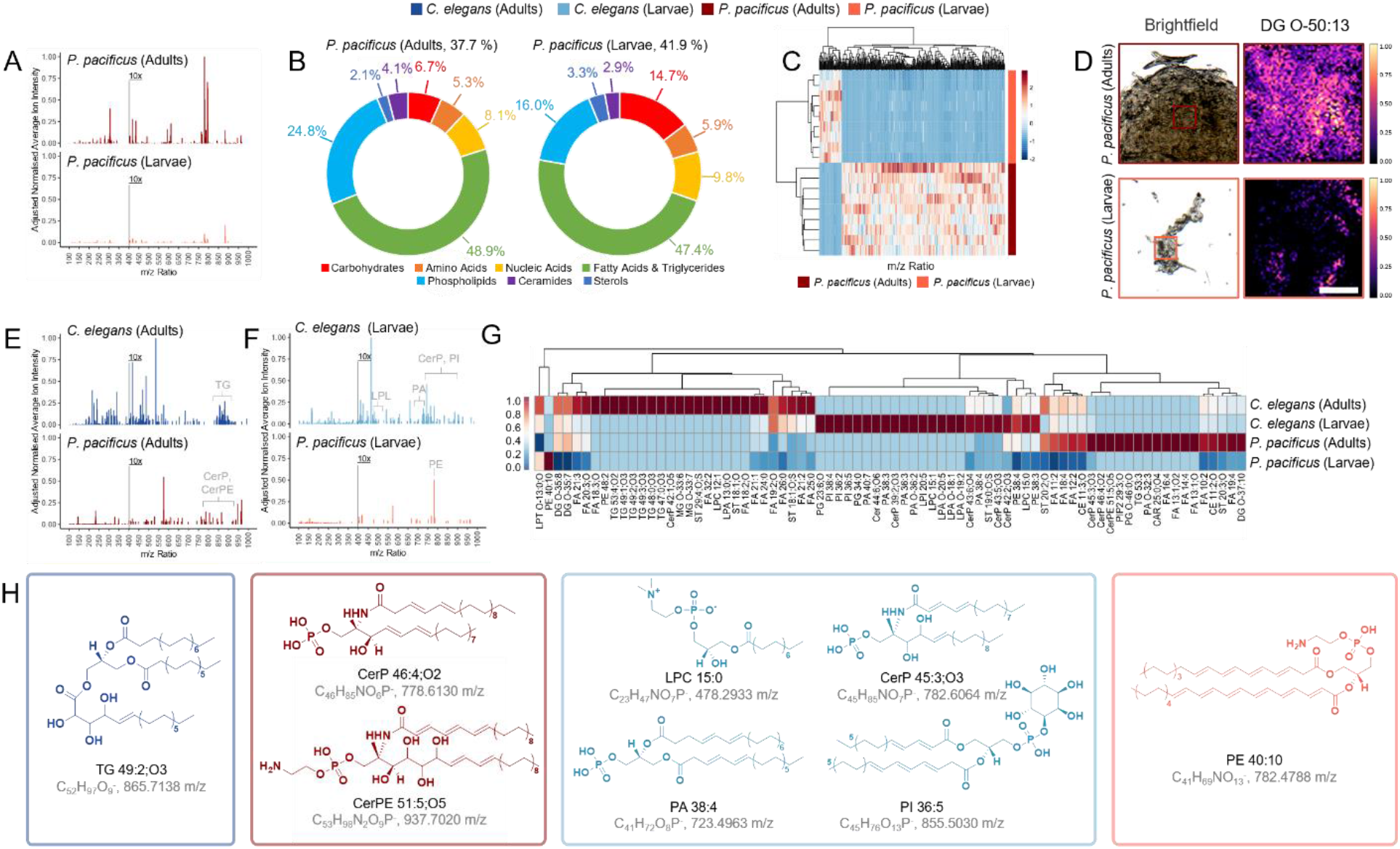
Surface chemistries are developmental stage dependant. **A** Averaged *P. pacificus* adults and larvae surface secondary ion mass spectra, normalised to maximum intensity across spectra, where intensity *m/z* >400 enhanced 10x for visibility. **B** Distribution of molecular assignments determined using chemical filtration (**Table S1**), as a percentage of total ions surveyed. **C** Hierarchical clustering heatmap for *P. pacificus* adults and larvae *m/z* ratio intensity, for PC1 loadings with a standard deviation greater than the mean. **D** Representative normalised intensity maps of *P. pacificus* adults and larvae surface chemistry, scale = 100 µm. Averaged surface secondary ion mass spectra exclusive to *C. elegans* and *P. pacificus* (**E** adults and **F** larvae). **G** Hierarchical clustering heatmaps of distribution of exclusive chemistries for *C. elegans (*adults & larvae) and *P. pacificus* (adults & larvae), where phylogeny indicates potential shared regulation of exclusive chemicals and their relative intensity on nematode surfaces. **H** Putative structures of significantly different chemistries on *C. elegans* and *P. pacificus* surfaces (*P*<0.001 by Student’s t-test, n=9), present in LIPIDS MAPS (M-H, < 2ppm).

The OrbiSIMS spectra for both *P. pacificus* adults and larvae were again dominated by surface-anchored lipids (**Fig. 2B**). Collectively, the total lipid components contributed to 79.96% and 69.61% of the OrbiSIMS spectra for adults and larvae, respectively. Both PCA (**Fig. S5**) and clustering (**Fig. 2C**) effectively differentiated the developmental stages of *P. pacificus*, revealing a reduction in the number of distinct mass ions characterising *P. pacificus* larvae compared to those observed in *C. elegans*. Molecular assignments and putative chemical structures reveal *P. pacificus* adults possess significantly more complex surface chemistries than larvae (**Fig. S5**). The predominant constituents are diglycerides (*e*.*g*., DG O-50:13), sterols (*e*.*g*., ST 20:0;O2), and a mix of unsaturated (*e*.*g*., FA 12:2) and saturated fatty acids (FA 20:0, FA 22:0, and FA 24:0). Representative normalised intensity maps of DG O-50:13, showcase the most pronounced relative intensity differences (**Fig. 3D**). This emphasises the observed variations in surface chemistry between P. pacificus adults and larvae and further delineates the developmental distinctions. These observations reveal stage-dependent chemical signals are also evident in other evolutionary diverse nematode species.

**Figure 3.**
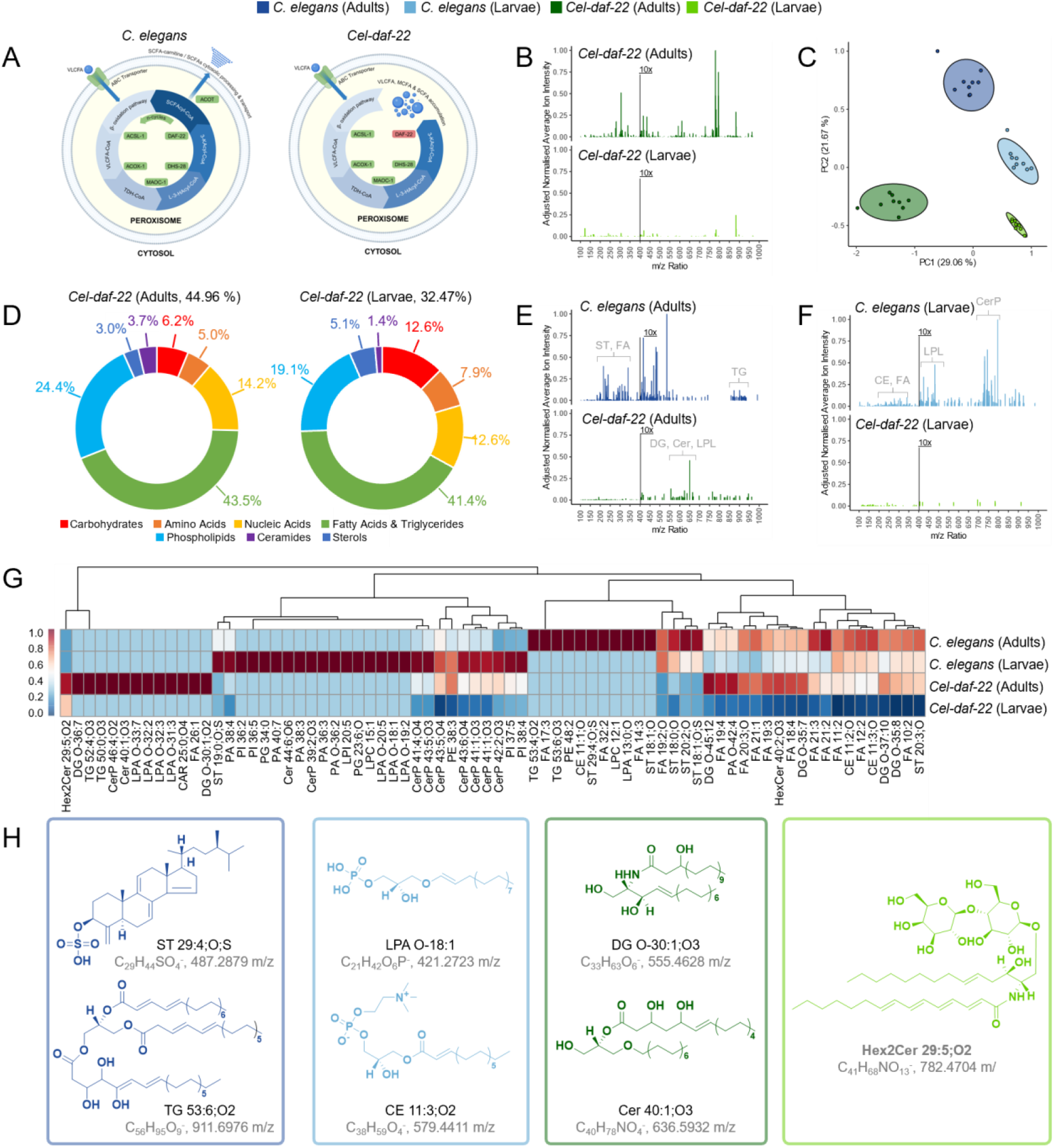
*C. elegans* surface profile is dependent on *daf-22*. **A** Schematic of peroxisomal β-oxidation pathway in wildtype and daf-22 mutation, restricting VLCFA, MCFA and SCFA processing and transport (created with BioRender.com). **B** Averaged *Cel-daf-22* adults and larvae surface secondary ion mass spectra, normalised to maximum intensity across spectra, where intensity *m/z* >400 enhanced 10x for visibility. **C** PCA PC1 & PC2 scores plot for *C. elegans* wildtype and *Cel-daf-22* developmental nematode stages. **D** Distribution of molecular assignments determined using chemical filtration (**Table S1**), as a percentage of total ions surveyed. Averaged surface secondary ion mass spectra exclusive to *C. elegans* and *Cel-daf-22 (***E** adults and **F** larvae). **G** Hierarchical clustering heatmaps of distribution of exclusive chemistries for *C. elegans* (adults & larvae) and *Cel-daf-22* (adults & larvae), where phylogeny indicates potential shared regulation of exclusive chemicals and their relative intensity on nematode surfaces. **H** Putative structures of significantly different chemistries on *C. elegans* wildtype and *daf-22* mutant surfaces (*P*<0.001 by Student’s t-test, n=9), present in LIPIDS MAPS (M-H, < 2ppm).*Acronyms:* ABC: ATP-binding cassette, ACOT: Acyl-CoA thioesterase, ACOX-1: Acyl-CoA oxidase 1, ACSL-1: Acyl-CoA synthetase long-chain family member 1, DAF-22: 3-ketoacyl-CoA thiolase, DHS-28: 3-hydroxyacyl-CoA dehydratase, L-3-HA-CoA: L-3-Hydroxyacyl-CoA, MAOC-1: Enoyl-CoA hydratase, CFA: Medium-chain fatty acids, SCFA: Short-chain fatty acids, TDH-CoA: Thiolase-CoA, VLCFA: Very long-chain fatty acids, 3-KA-CoA: 3-Ketoacyl-CoA.

### Species-specific surface chemistries

Having successfully identified the surface chemistries across developmental stages in two evolutionarily divergent nematodes, we next investigated species-specific differences in surface composition. The distribution of molecular assignments revealed significant variations in lipid composition between *C. elegans* and *P. pacificus* (**Fig. 1C, Fig. 2B**). In particular, the total lipid composition in *C. elegans* was consistently higher, with adults and larvae showing 3.75% and 14.84% greater lipid assignments, respectively, compared to *P. pacificus*. Specifically, the phospholipid composition in *C. elegans* larvae was 13.63% greater than in *P. pacificus* larvae, indicating species-specific and developmental stage lipid adaptations. These differences were further evidenced by PCA (**Fig. S6**) and hierarchical clustering heatmaps (**Fig. S7**), which confirmed species-specific and developmentally distinct surface chemistries. This suggests that *C. elegans* maintains higher lipid levels across developmental stages, unlike *P. pacificus*, where lipid content was considerably reduced in larvae.

By studying lipid components exclusive to each developmental stage and species, we discovered a unique cluster of secondary ions in *C. elegans* adults, ranging from m/z 850-900 (**Fig. 2E**), putatively identified as triglycerides (e.g., TG 48:0;O3, TG 49:2;O3, and TG 49:1;O3, **Fig. S8**). Whereas *P. pacificus* adults present secondary ions identified as ceramide phosphates (e.g., CerP 46:4;O2) and phosphoethanolamine (e.g., CerPE 51:5;O5) in the higher mass ranges. *C. elegans* larvae surfaces, compared to *P. pacificus* larvae (**Fig. 2F**), were also rich in ceramide phosphates (e.g. CerP 43:5,O3, CerP 43:6;O4 and CerP 45:3;O3) as well as a mixture of phosphatidylinositol (e.g., PI 36:5), phosphatidic acids (e.g., PA 38:4) and lysophosphatidic and lysophosphatidylcholine (**Fig. S8**). *P. pacificus* larvae in general, were absent of surface-specific lipids, compared to *C. elegans* larvae, except for a secondary ion at m/z 782.4788, which was putatively identified as phosphatidylethanolamine (PE 40:10, **Fig. 2G, Fig. 2H, Fig. S8**). While the nematodes share common surface components and biochemical pathways, these analyses highlight species-specific adaptations across divergent evolutionary paths enhancing our understanding of ecological diversity.

### Surface chemistries are *daf-22* dependent

As surface-anchored lipids were prominent on the *C. elegans* cuticle surface, we explored the role of metabolic pathways in producing these chemistries, by analysing mutants in the peroxisomal β-oxidation pathway. This pathway is essential for the degradation of very long-chain fatty acids and the synthesis of short chain fatty acids and ascarosides ^28^, with the thiolase DAF-22 acting as the terminal enzyme (**Fig. 3A**) ^29 30^. Therefore, we examined *daf-22* mutants in *C. elegans* to assess their impact on surface chemistries across both adult and larval stages (**Fig. 3B**). 3D-OrbiSIMS analysis revealed that *daf-22* mutants lacked many surface chemistries present in both wildtype adults and larvae (**Fig. 1C**). PCA effectively differentiated between the wildtype and *daf-22* mutants, which is consistent with substantial alterations to the surface chemistries in *daf-22* animals (**Fig. 3C, Fig. S9**).

Analysis of the distribution of molecular assignments *Cel-daf-22* adults and larvae exhibited abundance of surface-anchored lipids, akin to wildtype strains (**Fig. 3D)**. The total lipid component for *Cel-daf-22* adults and larvae was reduced by 8.5% and 14.85% in adults and larvae as in the percentage of lipid components compared to *C. elegans* wildtypes. In order to identify the precise effects of the *daf-22* mutations on the *C. elegans* surface composition, chemistries were isolated that were exclusive to either adult *C. elegans* wildtype or *daf-22* mutants (**Fig. 3E&F**). Adult *daf-22* mutants exhibited an absence of higher mass complex surface chemistries (*m/z* 850-950), putatively identified in wildtypes as unsaturated triglycerides (e.g., TG 53:4;O2 and TG 53:6;O3, **Fig. S10**), as well as a reduction in the relative abundance of lower mass ions *m/z* <400 consisting of unsaturated sterols and fatty acids (**Fig. 3E)**. Instead, an accumulation of mass ions between *m/z* 550–650 were found, putatively identified as diglycerides (*e*.*g*., DG O-30:1;O2) and ceramides (*e*.*g*., Cer 40:1;O3), as well as unsaturated lysophosphopholipids (**Fig. 3G**). The larval OrbiSIMS surface composition of *C. elegans daf-22* mutants also differed in a substantial number of chemistries when compared to wildtype at all *m/z* ratios (**Fig. 3F**). In particular, there was an absence of ceramide phosphates between *m/z* 700-800, lysophosphopholipids between *m/z* 400-500, and cholesteryl ester and unsaturated fatty acids between *m/z* 200-350 (**Fig. 3G, Fig. S10**). The diglycosylceramide Hex2Cer 29:5;O2 was also putatively found to have higher relative intensity in *Cel-daf-22* adults and larvae (**Fig. 3H**). Complex glycosphingolipids have been shown to modulate signal transduction pathways and influence behaviour, and simpler diglycosylceramide may have a similar function ^31^. Therefore, the peroxisomal β-oxidation pathway is essential and serves as a fundamental mechanism required for the synthesis of surface-anchored lipids and the establishment of stage-specific surface profiles in *C. elegans* development.

Given the divergent chemistries observed on *C. elegans* and *P. pacificus* surfaces, we also investigated the importance and conservation of the peroxisomal β-oxidation pathway for establishing the surface composition of *P. pacificus* using 3D-OrbiSIMS. Specifically, *P. pacificus* possesses two *daf-22* homologs, *Ppa-daf-22*.*1* and *Ppa-daf-22*.*2* ^32^. Analysis of the distribution of molecular component highlighted negligible differences in total lipid composition (<1 %) for both *Ppa-daf-22*.*1/2* double mutant adults and larvae, compared to wildtypes (**Fig. S11**). However, *Ppa-daf-22*.*1/2* mutants exhibited a reduction in specific secondary ions and their relative intensity (**Fig. S12 & S13**), indicating that although the overall lipid composition remains largely unchanged, certain lipid species are affected. This contrasts with the findings observed on *C. elegans* surfaces, where substantial changes in overall lipid composition were identified in *daf-22* mutants. Our observations suggest that *P. pacificus* may have compensatory mechanisms that maintain overall surface lipid levels despite disruptions in the peroxisomal β-oxidation pathway. These findings highlight differences in the regulatory mechanisms of surface lipid composition between these species and provide insight into the evolutionary adaptations of lipid metabolism pathways in nematodes.

### Surface chemistries regulate behaviours

Finally, given the distinct surface chemical profiles observed between *C. elegans* and *P. pacificus*, we hypothesised that these surface chemistries may function as species-specific communication signals and may influence contact-dependent behaviours. *P. pacificus* use its phenotypically plastic teeth-like denticles to direct predatory behaviour towards other nematode species (**Fig. 4A, Supplementary Video S1**) as well as other *P. pacificus* con-specifics, resulting in highly cannibalistic interactions ^10, 33, 34^. Crucially, there appears to be little influence from any volatile secreted molecules on this behaviour, which is instead determined by nose contact of the predator with the cuticle surface of a potential prey ^35^. Therefore, to test whether the altered surface lipid chemistries in *Cel-daf-22* serve as communication regulators, we conducted well-established predation assays using *P. pacificus* adults (**Fig. 4B**) ^10, 34^.

**Figure 4.**
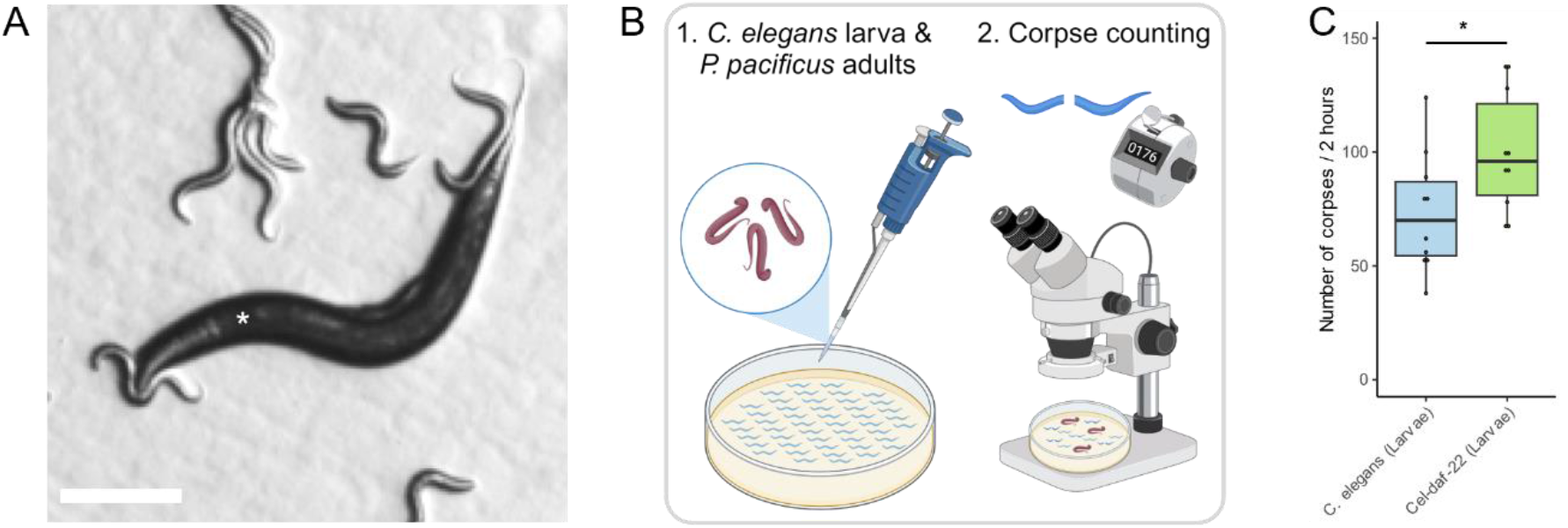
Surface chemistries regulating contact dependent predatory behaviour. **A** Representative images of P. pacificus wildtype (*) contact dependant predatory biting behaviour towards C. elegans wildtype larvae (scale = 100 µm). **B** Schematic of corpse assay of P. pacificus preying on C. elegans larvae (Created with BioRender.com). **c** Predatory behaviour of P. pacificus wildtype (adult) towards Cel-daf-22 larvae (*P<0.05 - Student’s t-test, n=10).

A significant difference in predation was observed between *C. elegans* wildtype and *Cel-daf-22* larvae (*P*=0.037, Student’s t-test, n=10 **Fig. 4C**). This indicates that the *daf-22* mutation leads to a shift and apparent reduction in surface lipid composition (**Fig. 3G**). This change increases susceptibility to predation, highlighting a potential protective contact-dependent function of these lipid. The increased predation of *Cel-daf-22* larvae likely stems from the altered surface lipid composition. The reduction in key structural lipids, such as fatty acids, sterols, and glycosphingolipids, may compromise the integrity of the cuticle, rendering the mutants more vulnerable to physical attack or alternatively, *daf-22* mutants may be more quickly identified as potential prey. (**Fig. 3G**). Furthermore, the alteration in surface composition, which was shown to produce novel groupings of lipids on mutants, could also disrupt membrane-dependent signalling pathways, potentially resulting in increased susceptibility to predation. These results confirm the multifaceted role of surface lipids in nematode survival, acting not only as structural components but also as mediators of inter-species communication and defence mechanisms.

## DISCUSSION

In this study, we have performed direct chemical analysis of the outermost 50 nm of the nematode surface using 3D-OrbiSIMS, we have generated an in-depth profile across two developmental stages and two evolutionary distinct species. This represents a significant advance on previous approaches, which utilised homogenate production, surface extraction steps ^17^, or relied on lower-resolution time-of-flight measurements ^20^. We reveal that the nematode surface profile is not a static entity, but instead is comprised of a complex, lipid-dominated landscape, which is dynamically modified during the organism’s development. Moreover, the lipid abundance and their hydrophobic properties across both species likely serve as a protective barrier against desiccation, enhancing our understanding of their essential roles beyond simple structural components. The *C. elegans* adult surface is characterised by the prevalence of complex lipid molecules, including longer-chain fatty acids and triglycerides, which are much less common on the larvae. These changes correlate with the maturing nematode metabolism ^36^, resulting in developmental stage specific signals that may be important for distinct population or environmental interactions. Furthermore, by comparing the surface composition across two distinct lineages of free-living nematodes, we also found the surface chemistry is species-specific indicating its importance as an evolutionary adaptive trait. While triglycerides dominate the surface of adult *C. elegans*, the surface of *P. pacificus* is instead comprised of ceramide phosphates, phosphatidylinositol and phosphatidic acids. In addition, the larval surface between species is also strikingly different, with the *P. pacificus* larval surface featuring fewer lipids contributing to its more naïve profile. This may represent a specific adaptation in this species, which could result in a concealed surface that may help avoid detection by predators.

The predatory behaviour observed in *P. pacificus* highlights the importance of species-specific recognition, likely mediated by surface lipid composition. The increased susceptibility of *C. elegans daf-22* mutants to predation underscores the protective role of an intact lipid profile. The *daf-22* mutation disrupts this profile, potentially impairing both the physical barrier function of the cuticle and the signalling pathways involved in predator avoidance. The surface hydrophobicity and signalling capabilities observed in nematodes parallel those seen in insects, where surface lipids not only prevent desiccation but also mediate communication regarding sex, age, reproductive status, and kinship ^37^.

Therefore, our studies not only highlight the complexity and dynamism of nematode surface chemistry but also reveal specific adaptations linked to their development and evolution. The use of advanced surface-sensitive mass spectrometry technology will enhance our understanding of surface lipid profiles but also paves the way for exploring subsurface chemistries and other complex behavioural interactions. For example, mapping complete metabolic processes across entire organisms could improve our understanding of host-parasitic nematode interactions. This may contribute to the development of novel strategies to overcome parasitic infections, thereby advancing public health interventions. We anticipate that expanding these approaches to a broader range of nematode species and environmental conditions, will further elucidate the evolutionary significance of surface chemistries and their role in ecological interactions.

## MATERIALS AND METHODS

### Nematode culture, sample preparation and behavioural assays

#### Nematode culture

All nematodes used were maintained on standard NGM plates on a diet of *Escherichia coli* OP50.

#### Nematode strains utilised

The following strains of *C. elegans* and *P. pacificus* were used in this study. The *C. elegans* strains included N2, the wildtype, and DR476, which has the *daf-22* (m130) mutation. For *P. pacificus*, the strains used were PS312, the wildtype, RS2770 with *daf-22*.*1* (tu489) and *daf-22*.*2* (tu504) mutations.

#### Nematode sample preparation

Nematodes were maintained on nematode growth medium (NGM) agar and *E. coli* (OP50) at 20 °C. Synchronized growth cycles were prepared by harvesting eggs from gravid females ^38^. Samples for analysis were gathered post synchronisation and after larval hatching from NMG agar plates (3 x 10 cm plates) using M9 buffer (10 mL, stored at 4 °C). Suspended nematodes were washed sequentially using centrifugation M9 (10 mL, 1500 rpm, 1 min x3), ammonium formate (150 mmol, 4 °C, 1500 rpm, 1 min x3) and ultra-pure water (18.2 MΩ, 1500 rpm, 1 min x 3) discarding the supernatant. After the final ultra-pure deionised water wash, nematodes were resuspended in ultrapure water and passed through two Nylon 20 µm filters (total volume 50 mL) to separate the adults from the larvae. Nematodes were pelleted with centrifugation; the worms were resuspended in a minimal volume of water. They were then deposited on indium tin oxide (ITO) slides (70-100 Ω)., excess water was dabbed off, and they were vacuum dried. Lastly, they were refrigerated (-80 °C) until analysis.

#### Corpse assay

Methods for corpse assay have been adapted from Lightfoot *et al*. ^39^ Briefly, Prey nematode cultures of *C. elegans* were grown on standard NGM plates supplemented with an *E. coli* OP50 lawn until freshly starved resulting in an abundance of young larvae. These plates were washed with M9 and passed through two 20 μm filters to isolate larvae which were collected in an Eppendorf tube. 1.0 μl of *C. elegans* larval pellet was transferred onto a 6 cm NGM unseeded plate. Five *P. pacificus* adult nematodes were transferred to plates of prey (larvae of wildtype *C. elegans* and *Cel-daf-22*). Assay plates were meticulously examined after 2 h for dead *C. elegans* larvae. These emptied corpses can be distinctly recognized by their immobility, coupled with evident morphological aberrations such as leaking internal organs or absent worm segments.

### Data acquisition

#### Scanning Electron Microscopy

Nematode specimens were initially synchronized to ensure uniform developmental stages. Post-synchronization, adult worms were washed multiple times M9 Buffer to remove any residual bacteria. The cleaned worms were fixed using 4% paraformaldehyde for 2 hours at room temperature. Following fixation, specimens were subjected to a series of dehydration steps using an increasing concentration of ethanol solutions (30%, 50%, 70%, 90%, and 100%) for 10 minutes each. The dehydrated samples were critical point dried to preserve their natural morphology and prevent shrinkage. Once dried, specimens were mounted onto carbon coated aluminium stubs. To improve conductivity and image quality, a thin layer of gold was sputter-coated onto the specimens. The coated nematode samples were then analysed under the SEM, focusing on the surface structures and morphology.

#### 3D-OrbiSIMS Data acquisition

Calibration of the Orbitrap analyser was performed using silver clusters, according to the protocol described by Passarelli et al. ^21^. The Bi_3_^+^ liquid metal ion gun and the ThermoFisher Tune software were executed for calibration. An Ar_3000_^+^ primary gas cluster ion beam (GCIB, 20 keV, 2 µm diameter, duty cycle set to 27.78% and current was 24 pA) was used for sample sputtering and ejection of secondary ions. The Q Exactive images were acquired using a random raster mode (field of view 300 × 300 µm, pixel size 5 µm, cycle time 200 μs, optimal analyser target -69.5 V). Argon gas flooding was in operation; to facilitate charge compensation. The pressure in the main sample chamber was also monitored and maintained (9.0 × 10^−7^ bar). The images were collected in negative polarity (mass range 75– *m/z* 1125) with constant injection time (500 ms) total ion dose per measurement (2.70 × 10^11^) and mass-resolving power (240,000 at *m/z* 200). Given the sputtering rate for organic materials, an ion dose of 3.00 × 10^14^ ions/cm^2^ was estimated to have analysed a sample depth of approximately 50 nm.

### Data extraction, management and filtration

#### Data extraction as well as noise identification and elimination

For each sample type (n=13) and region of interest (100 µm^2^), data was normalised to total ion count in SurfaceLab (version 7). Data was extracted and filtered by maximising data in each ROI using *C. elegans* wildtype adult as a reference. Through evaluation of the minimum intensity across the spectra, noise was determined as intensity counts less than 5×10^-6^ au, which identified ∼ 2500 secondary mass ions per ROI. The minimum noise (<5×10^-6^ au) and number of peaks (2500 secondary mass ions) criterion was translated to all additional data sets. Peak lists were combined to facilitate data comparison using a mass accuracy of 5 ppm. To eliminate noise generation during merged peak list production, data were surveyed to identify and eliminate newly generated noise with average noise intensity less than 5×10^-6^. The master peak list was composed of 11 different samples (adults & larvae for *C. elegans* wildtype, *Cel-daf-22, P. pacificus, Ppa-daf-22*.*1/2* as well as *E. coli*, agar and ITO Glass) each with 9 replicates (n=9, 100 µm^2^) and 9563 different peaks producing >1.1 million data points.

#### Mass Spectra Data Management and Visualization

The master peak list was searched to facilitate comparative analyses. Peak lists for comparative datasets were generated by combining data and removing data associated with *E. coli*, Agar and ITO, ensuring a <5 ppm mass accuracy, whilst applying an aggressive baseline threshold of 5×10^-6^ au for ions normalised to total ion count across each data set. These multivariate analyses were executed in R, utilising R Studio. Data were visualised as averaged secondary ion mass spectra, derived from ROI (n=9). The maximum filtered intensity was normalised and for data exceeding 400 *m/z*, a 10x multiplier was applied to facilitate data interpretation.

#### Distribution of molecular assignments

The distribution of molecular assignments was conducted using Secondary Ion Mass Spectrometry Molecular Formula Prediction (SIMSMFP) ^40^ and search criterion defined as carbohydrates (C_1-7_H_2-12_O_1-6_) proteins & amino acids (C_1-8_H_2-17_O_1-3_N_1_S_0-1_), nucleic Acids (C_1-8_H_2-9_O_1-6_N_1-3_P_1_), fatty Acids & triglycerides (C_2-50_H_4-90_O_1-4_), Phospholipids (C_4-40_H_8-80_O_4-8_N_1_P_1-2_) and sterols (C_10-25_H_9-44_O_1_), with mass deviations < 5 ppm. Instances where direct comparison of biological components between samples was required (*e*.*g*., *daf-22* mutant lipid component analysis) SIMSMFP was conducted to identify molecular components (*e*.*g*., fatty acids, triglycerides and phospholipids) which were then subtracted between organisms to < 5 ppm mass accuracy to determine differentiating mass ions.

#### Data analysis and packages

A range of R packages were used, each serving a specific purpose. SurfaceLab (Version 7) was used for data acquisition and manipulation, while R Studio (2022.07.02+576) provided an integrated development environment for R, the programming language version 4.2.2. The ‘dendextend’ package (version 1.17.1) aided in visualising and comparing hierarchical clustering trees. Data frame tools were handled by ‘dplyr’ (version 1.1.2), and ‘factoextra’ (version 1.0.7) and ‘FactoMineR’ (version 2.8) were used for visualization and extraction in multivariate analysis. The study also employed ‘forcats’ (version 1.0.0) for categorical variable tools, ‘ggforce’ (version 0.4.1) to accelerate ‘ggplot2’ (version 3.4.3), and ‘ggrepel’ (version 0.9.3) for positioning non-overlapping labels in ‘ggplot2’. Additional packages included ‘ggsignif’ (version 0.6.4), ‘ggtext’ (version 0.1.2), and ‘gplots’ (version 3.1.3) for plotting enhancements, ‘heatmaply’ (version 1.4.2) for interactive cluster heat maps, and ‘lattice’ (version 0.20-45) for trellis graphics. The ‘pheatmap’ package (version 1.0.12) was used for heatmap production, and ‘plotly’ (version 4.10.2) for interactive web graphics. The study also utilized ‘readr’ (version 2.1.4) for reading rectangular data, ‘scales’ (version 1.2.1) for scaling and formatting axis labels, ‘scatterplot3d’ (version 0.3-44) for 3D scatter plots, ‘stringr’ (version 1.5.0) for string operations, ‘svglite’ (version 2.1.1) for SVG graphics, ‘tibble’ (version 3.2.1) as a modern data.frame variant, and ‘tidyr’ (version 1.3.0) for data tidying. The ‘tidyverse’ collection (version 2.0.0) provided a comprehensive suite of data science packages, while ‘viridis’ (version 0.6.3) and ‘viridisLite’ (version 0.4.2) were used for creating perceptually uniform colour maps.

### Statistical analyses

All statistical analyses were performed in R using packages.

#### Comparative peak analyses

Student’s T-test (two tailed, unequal variance) were performed to identify significantly different mass during binary comparisons of datasets. Standard deviation was calculated between ROI (n=9) and plotted as error on accompanying bar charts. Significantly different data were categorized based on their p-value thresholds, such that * = P < 0.05 ** = *P* <0.01 and *** = *P* < 0.001.

#### Principal Component Analyses (PCA)

PCA was conducted on comparative data sets scaling and standardising and centring each variable so that they have standard deviation of 1 and a mean of 0, respectively. This step was important to ensure that all variables contribute equally to the principal components, eliminating any undue influence from variables. For 2D and 3D scores plots ellipse were used to visualise 95% confidence interval to provide a guide to consistency of data groupings.

#### Hierarchical clustering heat map analyses

Hierarchical clustering heatmap analysis, utilised a data reduction approach by selecting only mass ions with principal component loadings that were greater than one standard deviation from the mean. This data reduction was key to data visualisation, ensuring a focused analysis on the organisms and *m/z*-based separation of variables. Visualisation were optimised though application of heatmaps for columns of data comparing strains.

#### Putative Structural Molecular Assignments & Metabolic Pathway Analyses

Chemistries exhibiting significant differences were probed for potential structural assignments by searching the LIPIDMAPS ^25^ databases with a precision of ±0.0001 *m/z* (mass deviation of <2 ppm) and were normalized to nematode wildtype adults. Data were cross referenced with the Human Metabolome Database (precision ±0.0001 *m/z*, deviation of <2 ppm), to identify additional rational biologically derived chemistries not in LIPIDMAPS. The standard deviation served as error bars, and significance was determined using a Student’s T-test (P<0.001***, n=9). The identified putative chemical structures, with those found in all larvae shown in black and those deemed protective by PLS displayed in green, were further analysed.

## ASSOCIATED CONTENT

### Supporting Information

Supporting Data, including - Supporting Figures 1 to 13, Supporting Table S1, Supplementary Video S1 Caption, and Supporting Video 1 are available.

## AUTHOR INFORMATION

### Author Contributions

Conceptualization: VMC, JWL, Methodology: VMC, JWL, FH, AMK, Investigation: VMC, FH, AMK, Visualization: VMC, Funding acquisition: VMC, JWL, Project administration: VMC, Supervision: VMC, JWL, Writing – original draft: VMC, JWL, Writing – review & editing: VMC, JWL, DJS, MRA, FH, AMK

### Funding Sources

This work was funded by a Nottingham Research Fellowship, awarded by the University of Nottingham (VMC). This work was also funded by the grant number EP/P029868/1 (MRA, DJS). This work was also funded by Max Planck Society (JWL) and by the German Research Foundation (DFG) - project number 495445600 (JWL).

## Notes

### Data availability

All the data are publicly available via the Nottingham Research Data Management Repository at DOI: 10.17639/nott.7386

## Acknowledgment

We would like to thank Frankie Rawson (University of Nottingham, UK) as well as Monika Scholz and Desiree Götting (MPI Neurobiology of Behavior – caesar, Bonn) for discussion and critical reading of the manuscript. Additionally, we wish to thank the Sommer lab (MPI Biology, Tübingen) for providing *P. pacificus* strains. *C. elegans* strains were provided by the CGC, which is funded by NIH Office of Research Infrastructure Programs (P40 OD010440).

## Notes

### Competing Interest Statement

The authors have declared no competing interest.

### Summary of Updates

Title, abstract/summary, introduction, results, discussion, materials and methods, associated content, funding, and references.

## REFERENCES

(1) Agosta, W. C. Chemical communication: the language of pheromones; Henry Holt and Company, 1992.

(2) Ache, B. W.; Young, J. M. Olfaction: Diverse species, conserved principles. Neuron 2005, 48 (3), 417–430. DOI: 10.1016/j.neuron.2005.10.022.

(3) Butcher, R. A. Decoding chemical communication in nematodes. Natural Product Reports 2017, 34 (5), 472–477. DOI: 10.1039/c7np00007c.

(4) Faghih, N.; Bhar, S.; Zhou, Y.; Dar, A.; Mai, K.; Bailey, L.; Basso, K.; Butcher, R. A Large Family of Enzymes Responsible for the Modular Architecture of Nematode Pheromones. JOURNAL OF THE AMERICAN CHEMICAL SOCIETY 2020, 142, 13645–13650. DOI: 10.1021/jacs.0c04223.

(5) Ludewig, A. H.; Schroeder, F. C. Ascaroside signaling in C. elegans. WormBook. Jeong, P. Y.; Jung, M.; Yim, Y. H.; Kim, H.; Park, M.; Hong, E. M.; Lee, W.; Kim, Y. H.; Kim, K.; Paik, Y. K. Chemical structure and biological activity of the <i>Caenorhabditis elegans</i> dauer-inducing pheromone. Nature 2005, 433 (7025), 541–545. DOI: 10.1038/nature03201. Butcher, R. A.; Fujita, M.; Schroeder, F. C.; Clardy, J. Small-molecule pheromones that control dauer development in <i>Caenorhabditis elegans</i>. Nature Chemical Biology 2007, 3 (7), 420–422. DOI: 10.1038/nchembio.2007 Srinivasan, J.; Kaplan, F.; Ajredini, R.; Zachariah, C.; Alborn, H. T.; Teal, P. E. A.; Malik, R. U.; Edison, A. S.; Sternberg, P. W.; Schroeder, F. C. A blend of small molecules regulates both mating and development in <i>Caenorhabditis elegans</i>. Nature 2008, 454 (7208), 1115–U1146. DOI: 10.1038/nature07168. Butcher, R. A.; Ragains, J. R.; Kim, E.; Clardy, J. A potent dauer pheromone component in <i>Caenorhabditis elegans</i> that acts synergistically with other components. Proceedings of the National Academy of Sciences of the United States of America 2008, 105 (38), 14288–14292. DOI: 10.1073/pnas.0806676105. Macosko, E. Z.; Pokala, N.; Feinberg, E. H.; Chalasani, S. H.; Butcher, R. A.; Clardy, J.; Bargmann, C. I. A hub-and-spoke circuit drives pheromone attraction and social behaviour in <i>C-elegans</i>. Nature 2009, 458 (7242), 1171–U1110. DOI: 10.1038/nature07886. Srinivasan, J.; von Reuss, S. H.; Bose, N.; Zaslaver, A.; Mahanti, P.; Ho, M. C.; O’Doherty, O. G.; Edison, A. S.; Sternberg, P. W.; Schroeder, F. C. A Modular Library of Small Molecule Signals Regulates Social Behaviors in <i>Caenorhabditis elegans</i>. Plos Biology 2012, 10 (1). DOI: 10.1371/journal.pbio.1001237. von Reuss, S. H.; Bose, N.; Srinivasan, J.; Yim, J. J.; Judkins, J. C.; Sternberg, P. W.; Schroeder, F. C. Comparative Metabolomics Reveals Biogenesis of Ascarosides, a Modular Library of Small-Molecule Signals in <i>C</i>. <i>elegans</i>. Journal of the American Chemical Society 2012, 134 (3), 1817–1824. DOI: 10.1021/ja210202y.

(6) Sandhu, A.; Badal, D.; Sheokand, R.; Tyagi, S.; Singh, V. Specific collagens maintain the cuticle permeability barrier in <i>Caenorhabditis elegans</i>. Genetics 2021, 217 (3). DOI: 10.1093/genetics/iyaa047.

(7) Erkut, C.; Penkov, S.; Khesbak, H.; Vorkel, D.; Verbavatz, J. M.; Fahmy, K.; Kurzchalia, T. V. Trehalose Renders the Dauer Larva of <i>Caenorhabditis elegans</i> Resistant to Extreme Desiccation. Current Biology 2011, 21 (15), 1331–1336. DOI: 10.1016/j.cub.2011.06.064.

(8) Couillault, C.; Ewbank, J. J. Diverse bacteria are pathogens of <i>Caenorhabditis elegans</i>. Infection and Immunity 2002, 70 (8), 4705–4707. DOI: 10.1128/iai.70.8.4705-4707.2002.

(9) Hsueh, Y. P.; Mahanti, P.; Schroeder, F. C.; Sternberg, P. W. Nematode-Trapping Fungi Eavesdrop on Nematode Pheromones. Current Biology 2013, 23 (1), 83–86. DOI: 10.1016/j.cub.2012.11.035.

(10) Wilecki, M.; Lightfoot, J. W.; Susoy, V.; Sommer, R. J. Predatory feeding behaviour in Pristionchus nematodes is dependent on phenotypic plasticity and induced by serotonin. Journal of Experimental Biology 2015, 218 (9), 1306-1313, Article. DOI: 10.1242/jeb.118620.

(11) Adams, J. R. G.; Pooranachithra, M.; Jyo, E. M.; Zheng, S. L.; Goncharov, A.; Crew, J. R.; Kramer, J. M.; Jin, Y. S.; Ernst, A. M.; Chisholm, A. D. Nanoscale patterning of collagens in C. elegans apical extracellular matrix. Nature Communications 2023, 14 (1). DOI: 10.1038/s41467-023-43058-9.

(12) Cox, G. N.; Kusch, M.; Edgar, R. S. Cuticle of Caenorhabditis elegans - Its Isolation and Partial Characterization Journal of Cell Biology 1981, 90 (1), 7–17. DOI: 10.1083/jcb.90.1.7.

(13) Cox, G. N.; Kusch, M.; Edgar, R. S. The cuticle of Caenorhabditis elegans. II. Stage-specific changes in ultrastructure and protein composition during postembryonic development. Journal of Cell Biology 1981, 90 (1), 7–17. DOI: 10.1083/jcb.90.1.7.

(14) Hendriks, G. J.; Gaidatzis, D.; Aeschimann, F.; Grosshans, H. Extensive Oscillatory Gene Expression during <i>C</i>. <i>elegans</i> Larval Development. Molecular Cell 2014, 53 (3), 380–392. DOI: 10.1016/j.molcel.2013.12.013.

(15) Luo, J.; Bainbridge, C.; Miller, R.; Barrios, A.; Portman, D. C. elegans males optimize mate-preference decisions via sex-specific responses to multimodal sensory cues. CURRENT BIOLOGY 2024, 34. DOI: 10.1016/j.cub.2024.02.036 Weng, J.W.; Park, H.; Valotteau, C.; Chen, R.T.; Essmann, C.L.; Pujol, N.; Sternberg, P.W.; Chen, C.H. Body stiffness is a mechanical property that facilitates contact-mediated mate recognition in Caenorhabditis elegans. Current Biology 2023, 33 (17), 3585-+. DOI: 10.1016/j.cub.2023.07.020.

(16) Blaxter, M. L. Cuticle surface-proteins of wild-type and mutant Caenorhabditis elegans. Journal of Biological Chemistry 1993, 268 (9), 6600–6609. Palaima, E.; Leymarie, N.; Stroud, D.; Mizanur, R. M.; Hodgkin, J.; Gravato-Nobre, M. J.; Costello, C. E.; Cipollo, J. F. The <i>Caenorhabditis elegans</i> <i>bus</i>-<i>2</i> Mutant Reveals a New Class of <i>O</i>-Glycans Affecting Bacterial Resistance. Journal of Biological Chemistry 2010, 285 (23), 17662–17672. DOI: 10.1074/jbc.M109.065433. Gravato-Nobre, M. J.; Stroud, D.; O’Rourke, D.; Darby, C.; Hodgkin, J. Glycosylation Genes Expressed in Seam Cells Determine Complex Surface Properties and Bacterial Adhesion to the Cuticle of <i>Caenorhabditis elegans</i>. Genetics 2011, 187 (1), 141–155. DOI: 10.1534/genetics.110.122002. O’Rourke, D.; Gravato-Nobre, M. J.; Stroud, D.; Pritchett, E.; Barker, E.; Price, R. L.; Robinson, S. A.; Spiro, S.; Kuwabara, P.; Hodgkin, J. Isolation and molecular identification of nematode surface mutants with resistance to bacterial pathogens. G3-Genes Genomes Genetics 2023, 13 (5). DOI: 10.1093/g3journal/jkad056.

(17) Penkov, S.; Ogawa, A.; Schmidt, U.; Tate, D.; Zagoriy, V.; Boland, S.; Gruner, M.; Vorkel, D.; Verbavatz, J.-M.; Sommer, R. J.; et al. A wax ester promotes collective host finding in the nematode Pristionchus pacificus. Nature Chemical Biology 2014, 10 (4), 281-+. DOI: 10.1038/nchembio.1460.

(18) Menger, R. F.; Clendinen, C. S.; Searcy, L. A.; Edison, A. S.; Yost, R. A. MALDI Mass Spectrometric Imaging of the Nematode <i>Caenorhabditis elegans</i>. Current Metabolomics 2015, 3 (2), 130–137. DOI: 10.2174/2213235x03666150525223412.

(19) Chauhan, V. M.; Scurr, D. J.; Christie, T.; Telford, G.; Aylott, J. W.; Pritchard, D. I. The physicochemical fingerprint of Necator americanus. Plos Neglected Tropical Diseases 2017, 11 (12), 19, Article. DOI: 10.1371/journal.pntd.0005971.

(20) Geier, F. M.; Fearn, S.; Bundy, J. G.; McPhail, D. S. ToF-SIMS analysis of biomolecules in the model organism Caenorhabditis elegans. Surface and Interface Analysis 2013, 45 (1), 234–236. DOI: 10.1002/sia.5110.

(21) Passarelli, M. K.; Pirkl, A.; Moellers, R.; Grinfeld, D.; Kollmer, F.; Havelund, R.; Newman, C. F.; Marshall, P. S.; Arlinghaus, H.; Alexander, M. R.; et al. The 3D OrbiSIMS-label-free metabolic imaging with subcellular lateral resolution and high mass-resolving power. Nature Methods 2017, 14 (12), 1175–1190, Article. DOI: 10.1038/nmeth.4504.

(22) Kotowska, A. M.; Trindade, G. F.; Mendes, P. M.; Williams, P. M.; Aylott, J. W.; Shard, A. G.; Alexander, M. R.; Scurr, D. J. Protein identification by 3D OrbiSIMS to facilitate in situ imaging and depth profiling. Nature Communications 2020, 11 (1), 5832. DOI: 10.1038/s41467-020-19445-x. Starr, N. J.; Khan, M. H.; Edney, M. K.; Trindade, G. F.; Kern, S.; Pirkl, A.; Kleine-Boymann, M.; Elms, C.; O’Mahony, M. M.; Bell, M. Elucidating the molecular landscape of the stratum corneum. Proceedings of the National Academy of Sciences 2022, 119 (12), e2114380119.

(23) Bailey, J.; Havelund, R.; Shard, A. G.; Gilmore, I. S.; Alexander, M. R.; Sharp, J. S.; Scurr, D. J. 3D ToF-SIMS Imaging of Polymer Multi layer Films Using Argon Cluster Sputter Depth Profiling. Acs Applied Materials & Interfaces 2015, 7 (4), 2654–2659. DOI: 10.1021/am507663v.

(24) La Cavera, S.; Chauhan, V. M.; Hardiman, W.; Yao, M.; Fuentes-Domínguez, R.; Setchfield, K.; Abayzeed, S. A.; Pérez-Cota, F.; Smith, R. J.; Clark, M. Label-free Brillouin endo-microscopy for the quantitative 3D imaging of sub-micrometre biology. Communications Biology 2024, 7 (1), 451. DOI: 10.1038/s42003-024-06126-4.

(25) Sud, M.; Fahy, E.; Cotter, D.; Brown, A.; Dennis, E. A.; Glass, C. K.; Merrill, A. H.; Murphy, R. C.; Raetz, C. R. H.; Russell, D. W.; et al. LMSD: LIPID MAPS structure database. Nucleic Acids Research 2007, 35, D527-D532. DOI: 10.1093/nar/gkl838.

(26) Félix, M. A.; Duveau, F. Population dynamics and habitat sharing of natural populations of <i>Caenorhabditis elegans</i> and <i>C</i>. <i>briggsae</i>. Bmc Biology 2012, 10. DOI: 10.1186/1741-7007-10-59.

(27) Herrmann, M.; Mayer, W. E.; Sommer, R. J. Nematodes of the genus <i>Pristionchus</i> are closely associated with scarab beetles and the Colorado potato beetle in Western Europe. Zoology 2006, 109 (2), 96–108. DOI: 10.1016/j.zool.2006.03.001.

(28) Zhang, S. B. O.; Box, A. C.; Xu, N. Y.; Le Men, J.; Yu, J. Y.; Guo, F. L.; Trimble, R.; Mak, H. Y. Genetic and dietary regulation of lipid droplet expansion in <i>Caenorhabditis elegans</i>. Proceedings of the National Academy of Sciences of the United States of America 2010, 107 (10), 4640–4645. DOI: 10.1073/pnas.0912308107. Butcher, R. A.; Ragains, J. R.; Li, W. Q.; Ruvkun, G.; Clardy, J.; Mak, H. Y. Biosynthesis of the <i>Caenorhabditis elegans</i> dauer pheromone. Proceedings of the National Academy of Sciences of the United States of America 2009, 106 (6), 1875–1879. DOI: 10.1073/pnas.0810338106. Artyukhin, A. B.; Zhang, Y. K.; Akagi, A. E.; Panda, O.; Sternberg, P. W.; Schroeder, F. C. Metabolomic “Dark Matter” Dependent on Peroxisomal beta-Oxidation in Caenorhabditis elegans. Journal of the American Chemical Society 2018, 140 (8), 2841–2852. DOI: 10.1021/jacs.7b11811.

(29) Joo, H.; Yim, Y.; Jeong, P.; Jin, Y.; Lee, J.; Kim, H.; Jeong, S.; Chitwood, D.; Paik, Y. <i>Caenorhabditis elegans</i> utilizes dauer pheromone biosynthesis to dispose of toxic peroxisomal fatty acids for cellular homoeostasis. BIOCHEMICAL JOURNAL 2009, 422, 61–71. DOI: 10.1042/BJ20090513.

(30) Wang, Y.; Li, C.; Zhang, J.; Xu, X.; Fu, L.; Xu, J.; Zhu, H.; Hu, Y.; Li, C.; Wang, M.; et al. Polyunsaturated fatty acids promote the rapid fusion of lipid droplets in <i>Caenorhabditis elegans</i>. JOURNAL OF BIOLOGICAL CHEMISTRY 2022, 298. DOI: 10.1016/j.jbc.2022.102179.

(31) Hakomori, S. The glycosynapse. PROCEEDINGS OF THE NATIONAL ACADEMY OF SCIENCES OF THE UNITED STATES OF AMERICA 2002, 99, 225–232.

(32) Markov, G. V.; Meyer, J. M.; Panda, O.; Artyukhin, A. B.; Claassen, M.; Witte, H.; Schroeder, F. C.; Sommer, R. J. Functional Conservation and Divergence of <i>daf</i>-<i>22</i> Paralogs in <i>Pristionchus pacificus</i> Dauer Development. Molecular Biology and Evolution 2016, 33 (10), 2506–2514. DOI: 10.1093/molbev/msw090.

(33) Ragsdale, E. J.; Müller, M. R.; Rödelsperger, C.; Sommer, R. J. A Developmental Switch Coupled to the Evolution of Plasticity Acts through a Sulfatase. Cell 2013, 155 (4), 922–933. DOI: 10.1016/j.cell.2013.09.054. Quach, K. T.; Chalasani, S. H. Flexible reprogramming of Pristionchus pacificus motivation for attacking Caenorhabditis elegans in predator-prey competition. Current Biology 2022, 32 (8), 1675-+. DOI: 10.1016/j.cub.2022.02.033. Lightfoot, J. W.; Dardiry, M.; Kalirad, A.; Giaimo, S.; Eberhardt, G.; Witte, H.; Wilecki, M.; Rodelsperger, C.; Traulsen, A.; Sommer, R. J. Sex or cannibalism: Polyphenism and kin recognition control social action strategies in nematodes. Science Advances 2021, 7 (35). DOI: 10.1126/sciadv.abg8042. Hiramatsu, F.; Lightfoot, J. W. Kin-recognition and predation shape collective behaviors in the cannibalistic nematode <i>Pristionchus pacificus</i>. Plos Genetics 2023, 19 (12). DOI: 10.1371/journal.pgen.1011056.

(34) Lightfoot, J. W.; Wilecki, M.; Rodelsperger, C.; Moreno, E.; Susoy, V.; Witte, H.; Sommer, R. J. Small peptide-mediated self-recognition prevents cannibalism in predatory nematodes. Science 2019, 364 (6435), 86-+. DOI: 10.1126/science.aav9856.

(35) Moreno, E.; Lightfoot, J. W.; Lenuzzi, M.; Sommer, R. J. Cilia drive developmental plasticity and are essential for efficient prey detection in predatory nematodes. Proceedings of the Royal Society B-Biological Sciences 2019, 286 (1912). DOI: 10.1098/rspb.2019.1089.

(36) Gao, A. W.; Chatzispyrou, I. A.; Kamble, R.; Liu, Y. J.; Herzog, K.; Smith, R. L.; van Lenthe, H.; Vervaart, M. A. T.; van Cruchten, A.; Luyf, A. C.; et al. A sensitive mass spectrometry platform identifies metabolic changes of life history traits in C. elegans. Scientific Reports 2017, 7. DOI: 10.1038/s41598-017-02539-w.

(37) Ozaki, M.; Wada-Katsumata, A.; Fujikawa, K.; Iwasaki, M.; Yokohari, F.; Satoji, Y.; Nisimura, T.; Yamaoka, R. Ant nestmate and non-nestmate discrimination by a chemosensory sensillum. Science 2005, 309 (5732), 311–314. DOI: 10.1126/science.1105244.

(38) Chauhan, V. M.; Orsi, G.; Brown, A.; Pritchard, D. I.; Aylott, J. W. Mapping the Pharyngeal and Intestinal pH of Caenorhabditis elegans and Real-Time Luminal pH Oscillations Using Extended Dynamic Range pH-Sensitive Nanosensors. Acs Nano 2013, 7 (6), 5577-5587, Article. DOI: 10.1021/nn401856u.

(39) Lightfoot, J. W.; Wilecki, M.; Okumura, M.; Sommer, R. J. Assaying Predatory Feeding Behaviors in Pristionchus and Other Nematodes. Jove-Journal of Visualized Experiments 2016, (115). DOI: 10.3791/54404.

(40) Edney, M. K.; Kotowska, A. M.; Spanu, M.; Trindade, G. F.; Wilmot, E.; Reid, J.; Barker, J.; Aylott, J. W.; Shard, A. G.; Alexander, M. R.; et al. Molecular Formula Prediction for Chemical Filtering of 3D OrbiSIMS Datasets. Analytical Chemistry 2022, 94 (11), 4703–4711. DOI: 10.1021/acs.analchem.1c04898.

