## Supporting Information for "Surface lipids influence developmental and species-specific chemical signalling in nematodes"

**This PDF file includes:**

- Supplementary Figures 1 to 13
- Supplementary Table S1
- Supplementary Video S1 Caption

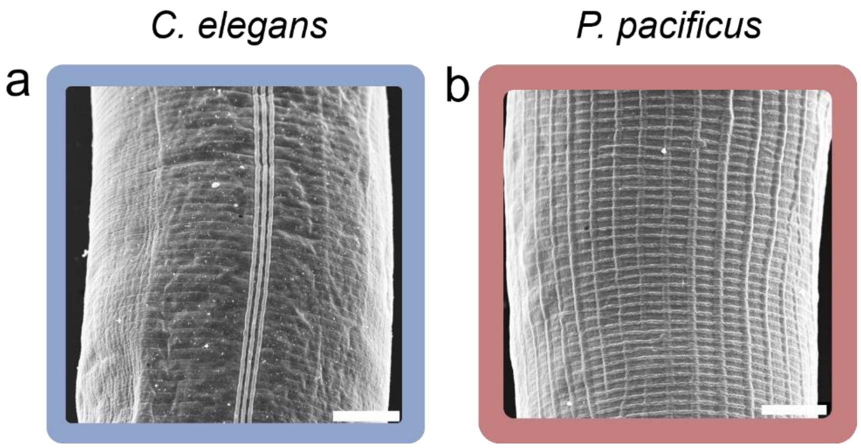


**Fig. S1 | *C. elegans* and *P. pacificus* rich topological features that generate the organism's external morphology .** SEM image of **a** C. elegans  and **b** *P. pacificus* surface cuticle, scale = 8 μm.

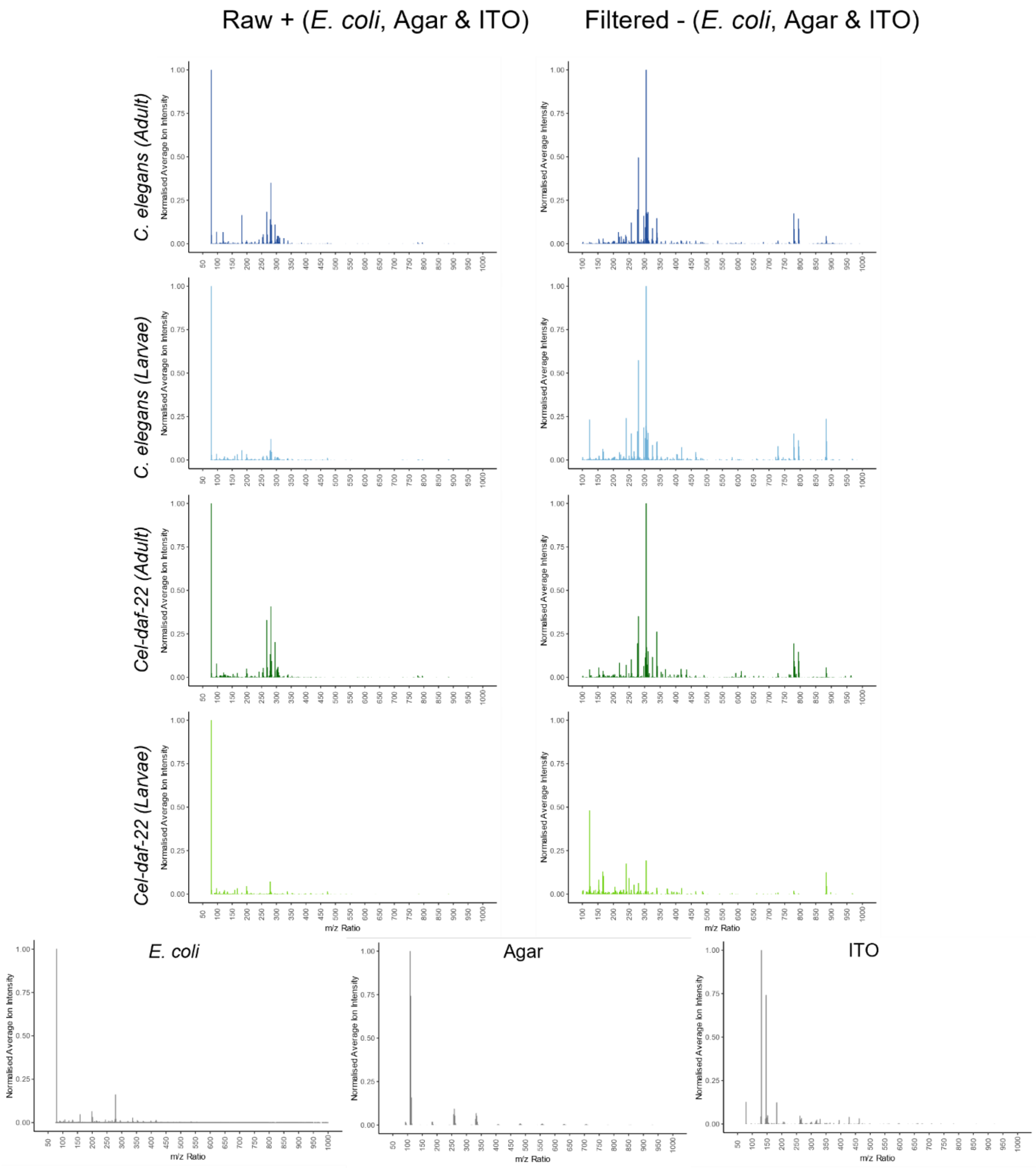


**Fig. S2 | *C. elegans* mass spectra optimisation.** Raw and filtered secondary ion mass spectra, eliminating *E.coli*, Agar and ITO, ions to 5 ppm.

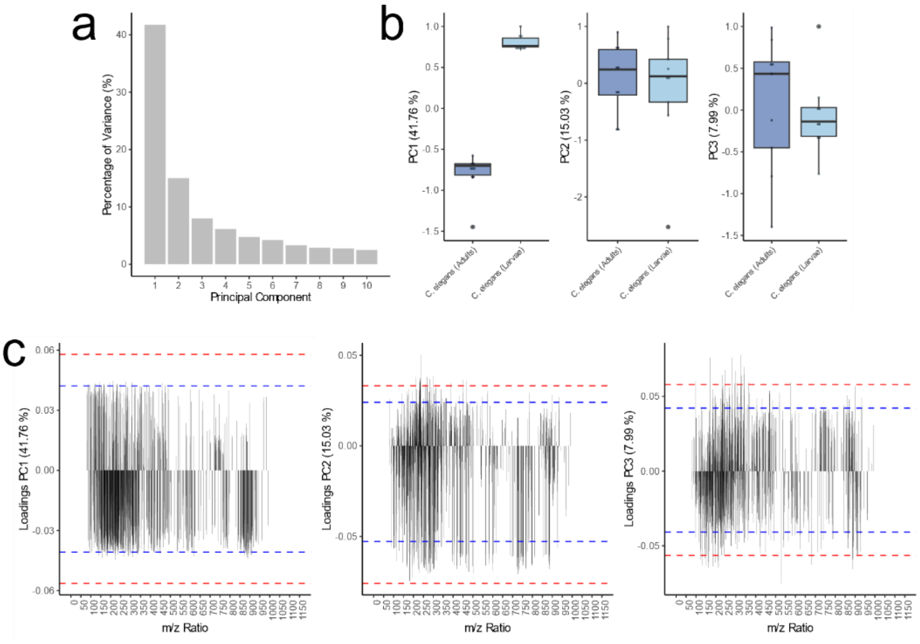


**Fig. S3 | Multivariate analysis of *C. elegans* adults and larvae. a** Percentage of variance on each principal component. **b** PC1, PC2 and PC3 b scores plot. **c** PC1, PC2 and PC3 loadings, where blue and red dashed lines indicate 1 and 2 standard deviations from the mean, respectively.

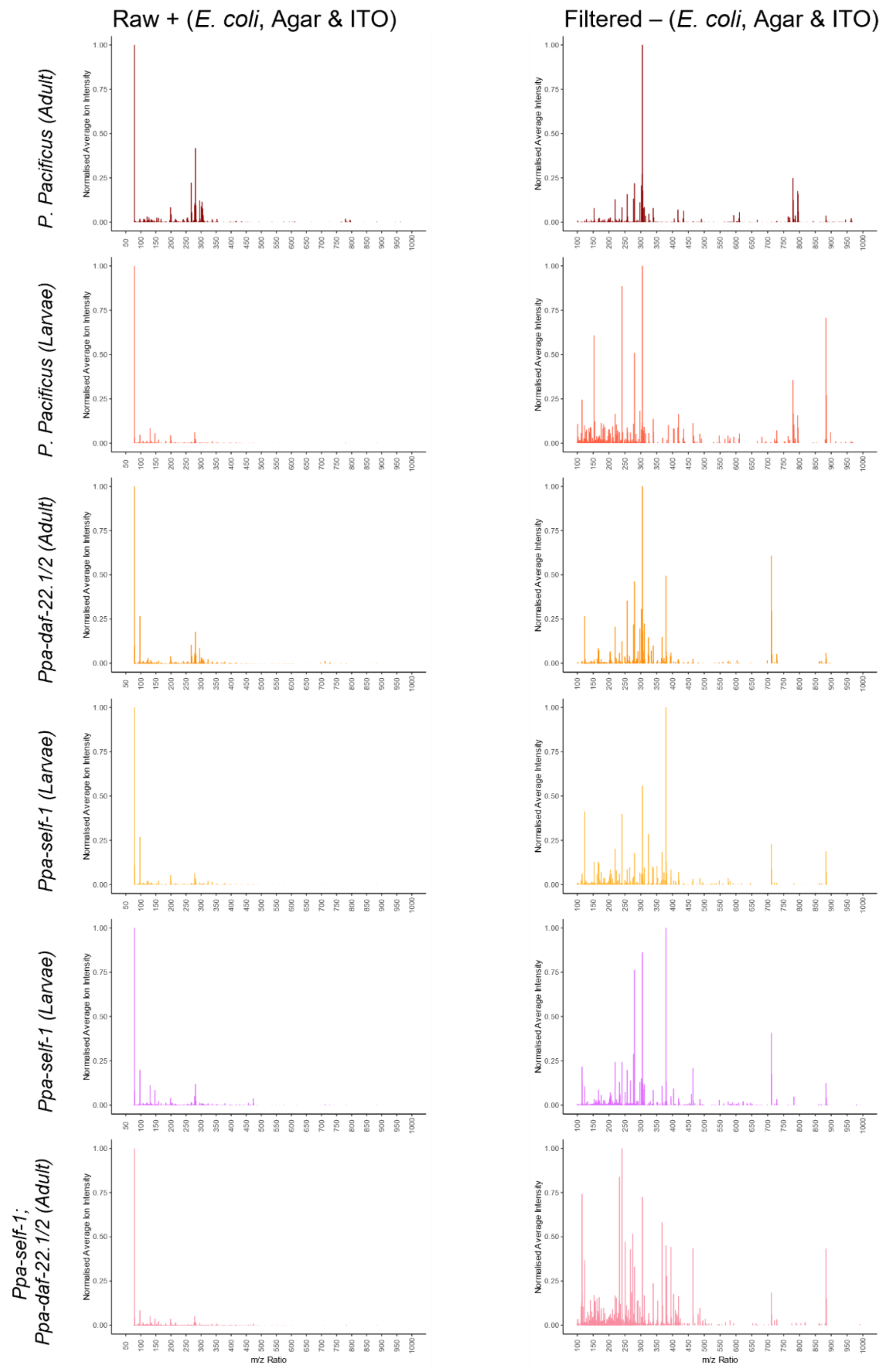


**Fig. S4 | *P. pacificus* mass spectra optimisation.** Raw and filtered secondary ion mass spectra, eliminating *E.coli*, Agar and ITO, ions to 5 ppm.

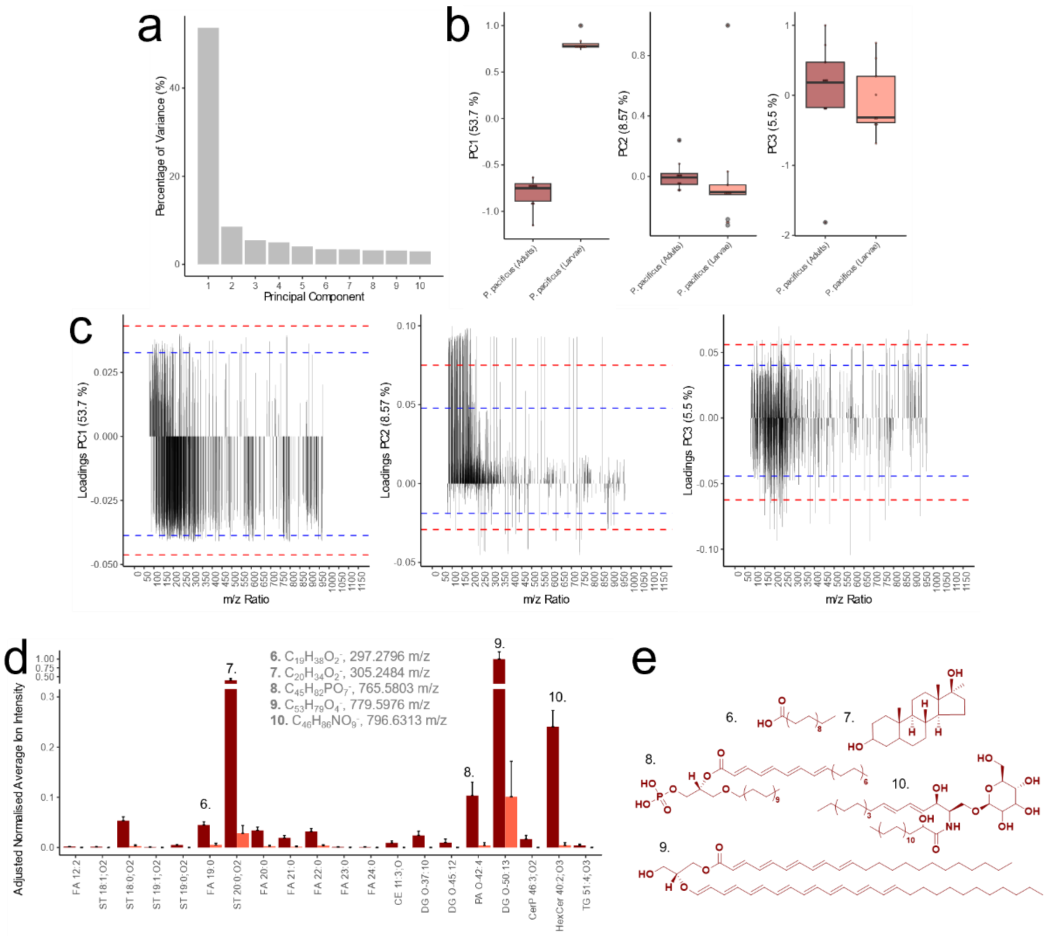


**Fig. S5 | Multivariate analysis of *P. pacificus* adults and larvae. a** Percentage of variance on each principal component. **b** PC2 and PC3 scores plot. **c** PC1, Pc2 and PC3 loadings, where blue and red dashed lines indicate 1 and 2 standard deviations from the mean, respectively. Significantly different chemistries on *P. pacificus* adults and larvae surfaces, where (*P*<0.001 by Student's t-test, n=9), present in LIPIDS MAPS (M-H, < 2 ppm) with putative **d** chemical assignments and **e** structures.

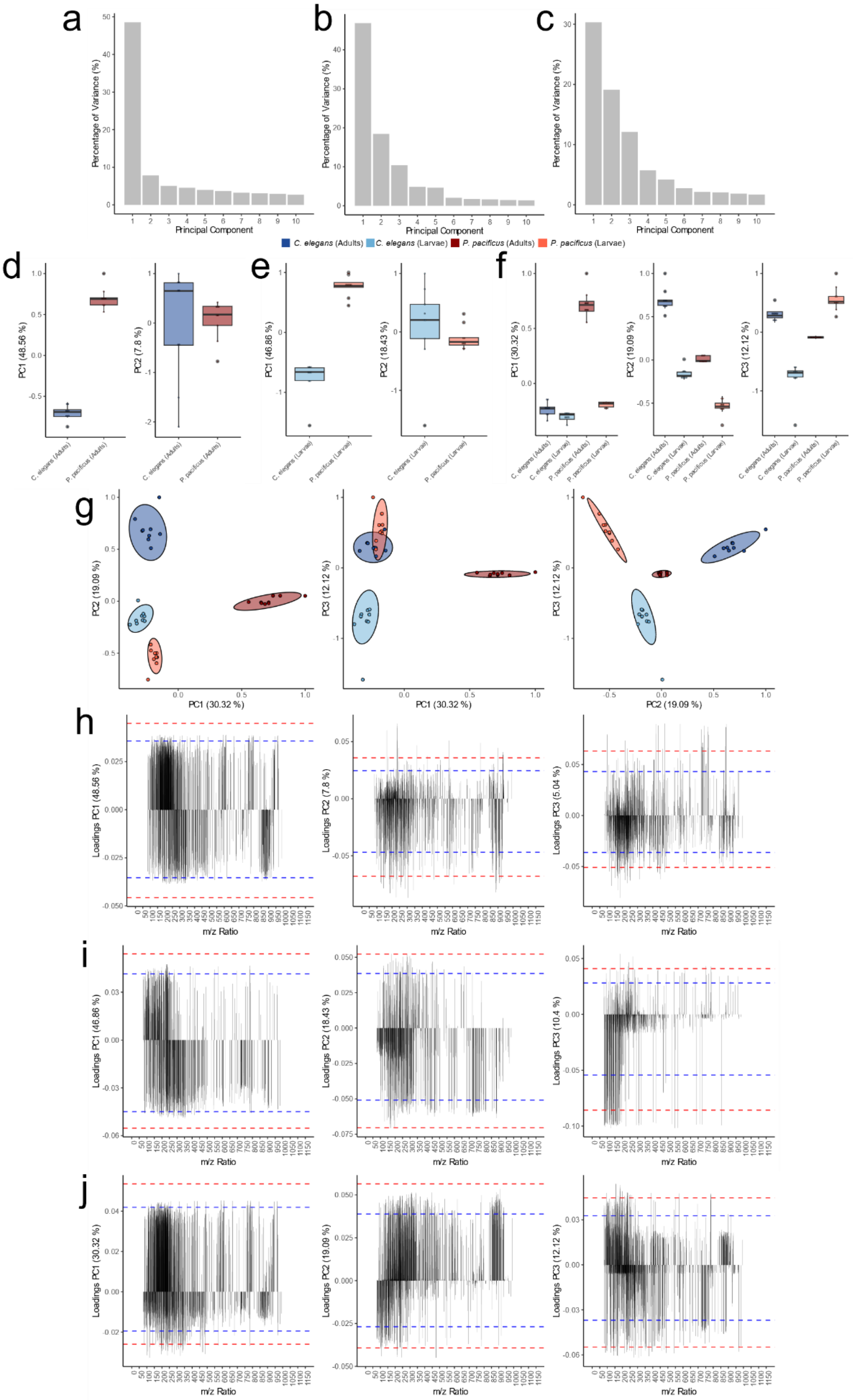


**Fig. S6 | Multivariate analysis of *C. elegans* and *P. pacificus* developmental stages. a** Percentage of variance on each principal component (**a** adults **b** larvae and **c** adults & larvae). PC1 and PC2 scores plot (**d** adult **e** larvae). **f** PC1, PC2 and PC3 *C. elegans* and *P. pacificus* developmental stages. **g** PC1&2, PC1&3 and PC2&3 scores biplots for *C. elegans* and *P. pacificus* developmental stages. PC1, PC2 and PC3 loadings plots , where blue and red dashed lines indicate 1 and 2 standard deviations from the mean, respectively (**h** adults **i** larvae and **j** adults & larvae).

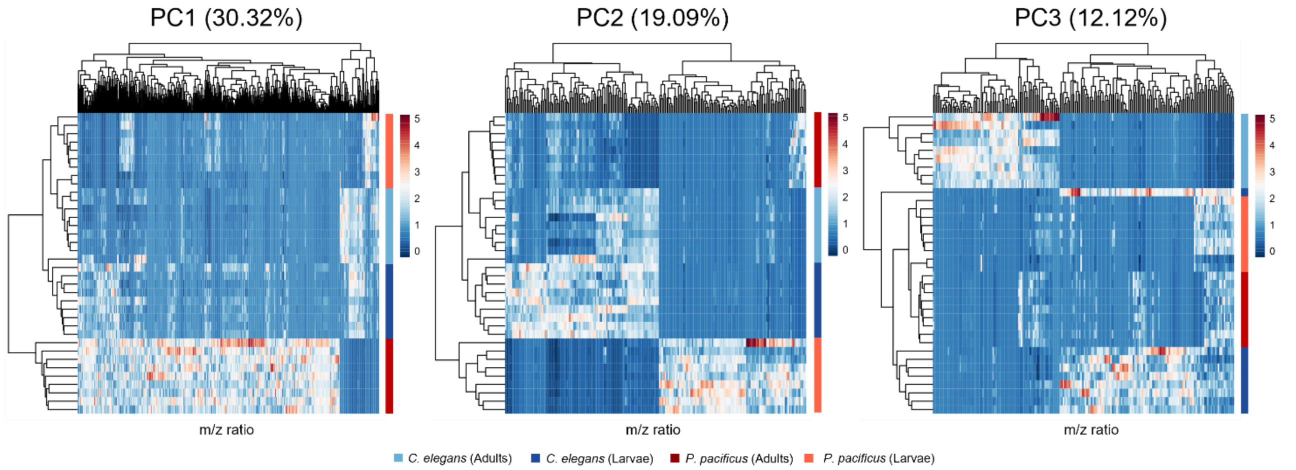


**Fig. S7 | Comparing *C. elegans* and *P. pacificus* hierarchical clustering heatmaps.** Hierarchical clustering heatmaps for subsets of data exhibiting loadings greater than 1 standard deviation from the mean using PC1-3 PCA analysis.

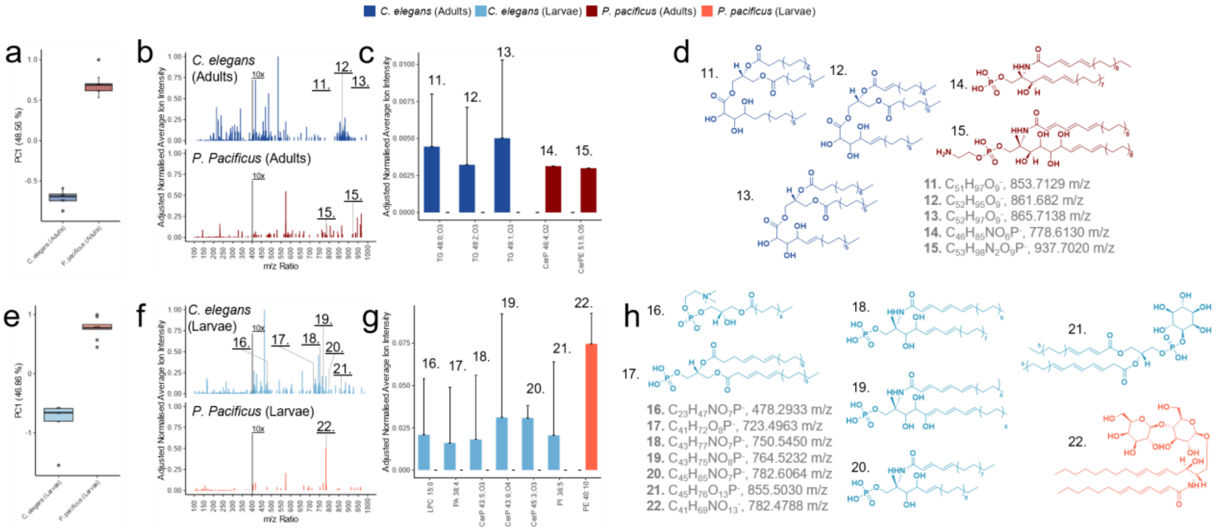


**Fig. S8 | Species-specific surface chemical contrast divergence.** PCA PC1 scores plot for *C. elegans* and *P. pacificus* (**a** adults and **e** larvae). Averaged surface secondary ion mass spectra exclusive to *C. elegans* and *P. pacificus* (**b** adults and **f** larvae) . Putative chemical assignments (**c** adults and **g** larvae) and structures (**d** adults and **h** larvae) on *C. elegans* and *P. pacificus* surfaces (*P*<0.001 by Student's t-test, n=9), present in LIPIDS MAPS (M-H, < 2ppm).

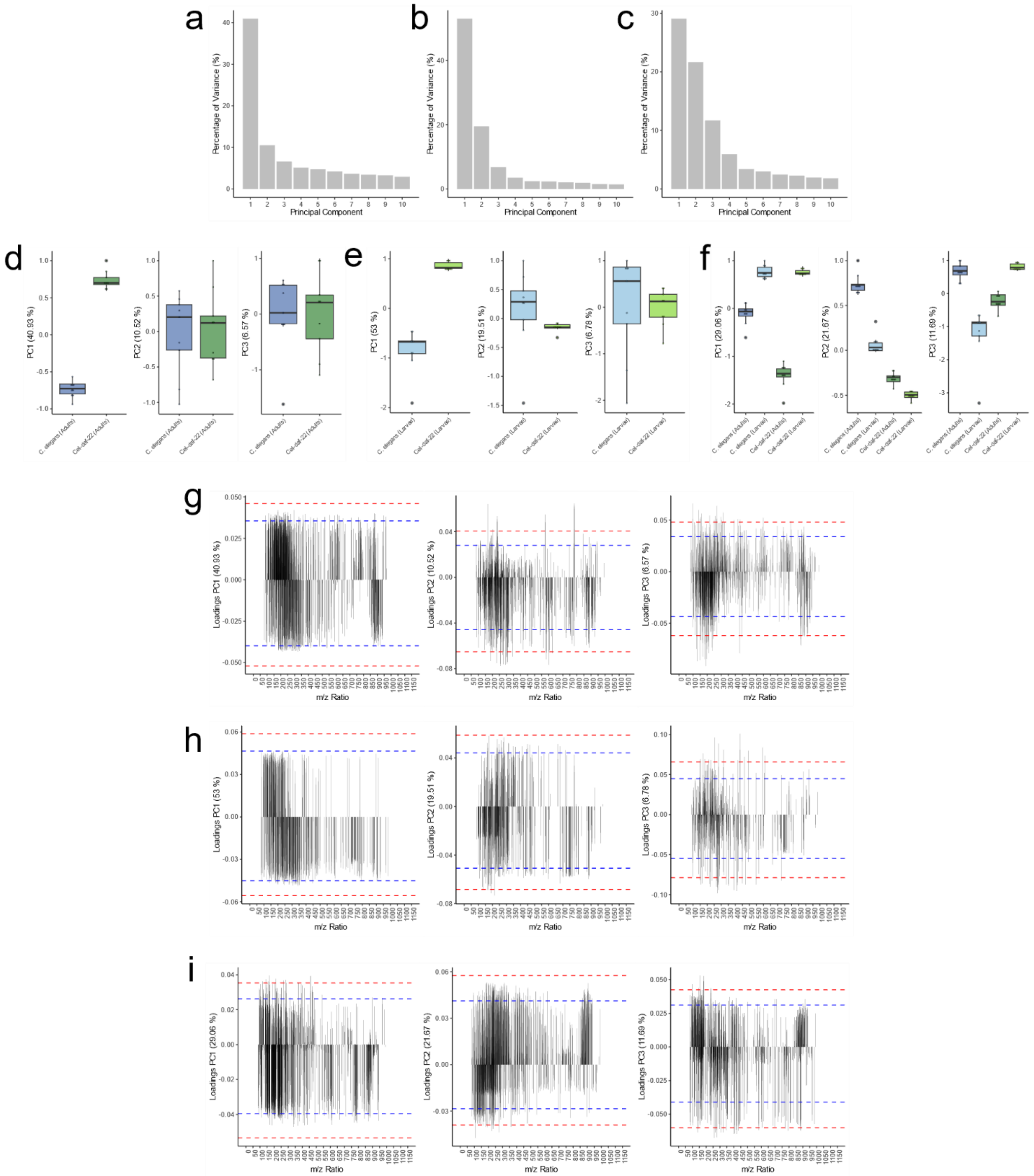


**Fig. S9 | Multivariate analysis of *C. elegans* wildtype and *daf-22* for each developmental stage. a** Percentage of variance on each principal component (**a** adults, **b** larvae and **c** adults & larvae). PC1, PC2 and PC3 scores plot (**d** adults and **e** larvae). **f** PC1, PC2 and PC3 wildtype and daf-22 developmental stages scores plot. PC1, PC2 and PC3 loadings, where blue and red dashed lines indicate 1 and 2 standard deviations from the mean, respectively (**g** adults, **h** larvae and **i** adults & larvae)

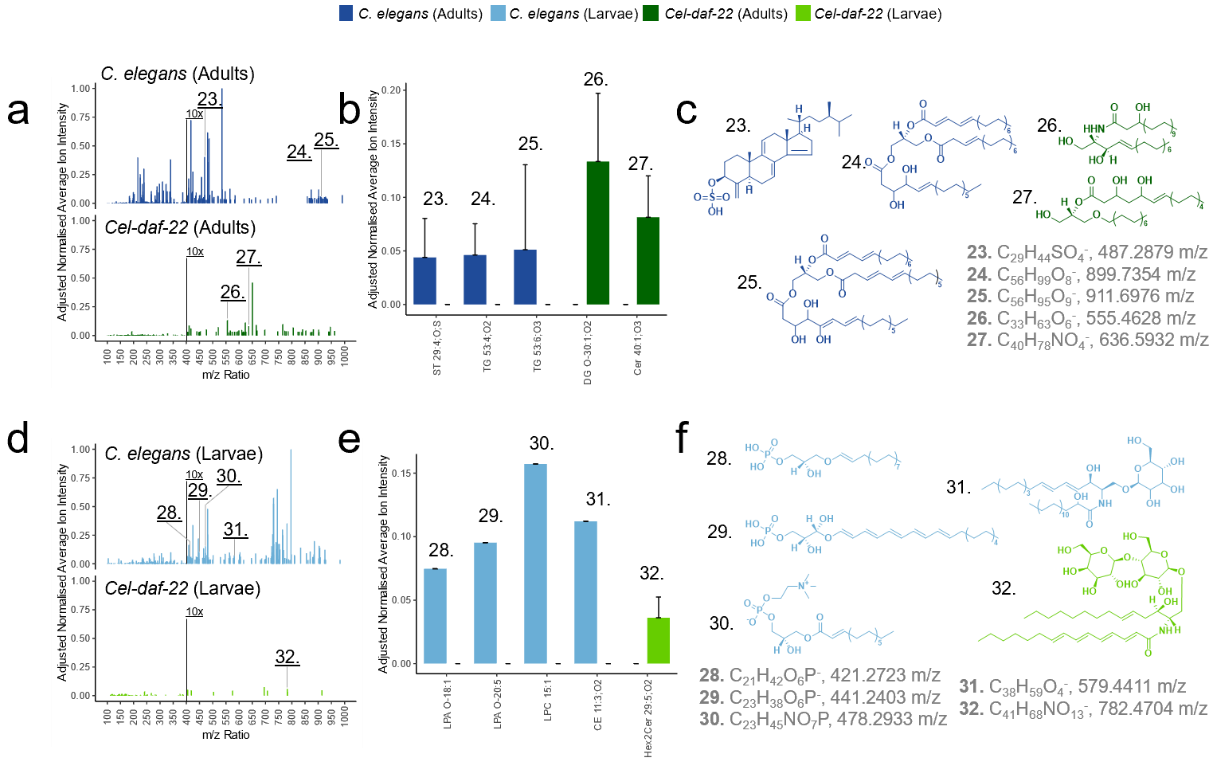


**Fig S10. Putative *C. elegans* wild-type and *daf-22* mutant chemical assignments and structures** Averaged surface secondary ion mass spectra exclusive to *C. elegans* and *Cel-daf-22* (**a** adults and **d** larvae). Putative chemical assignments (**b** adults and **e** larvae) and structures (**c** adults and **f** larvae) on *C. elegans* and *Cel-daf-22* surfaces ($P$<0.001 by Student's t-test, n=9), present in LIPIDS MAPS (M-H, < 2ppm).

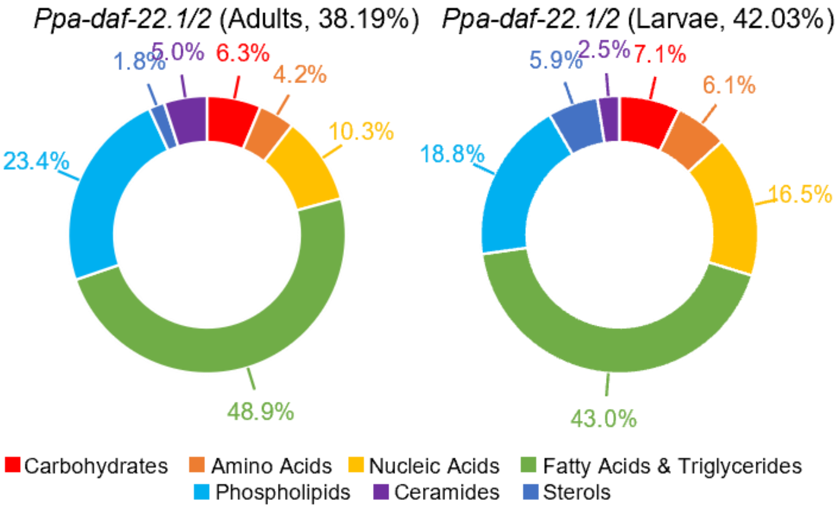


**Fig. S11 | *P. pacificus* surface profile is dependent on *daf-22*.** Distribution of molecular assignments determined using chemical filtration (**Table S1**), as a percentage of total ions surveyed for *Ppa-daf-22.1/2* adults and larvae surfaces.

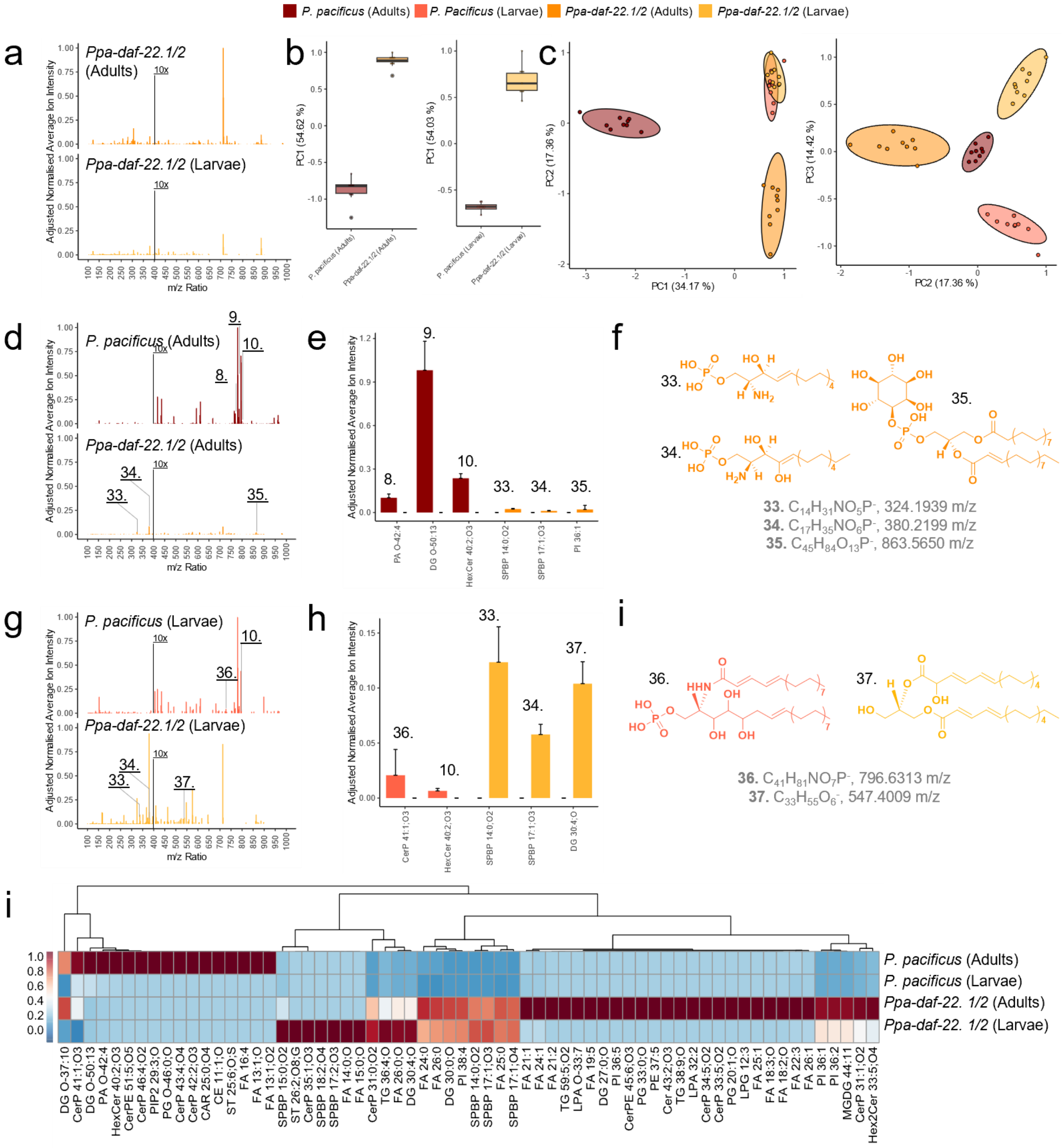

**Fig. S12 | *P. pacificus* surface profile is dependent on *daf-22*. a** Averaged *Ppa-daf-22.1/2* adults and larvae surface secondary ion mass spectra, normalised to maximum intensity across spectra, where intensity *m/z* >400 enhanced 10x for visibility. PCA **b** PC1 and **c** PC1 & PC2 and PC2 & PC3 scores plots for *P. pacificus* and *Ppa-daf-22.1/2* developmental nematode stages. Averaged surface secondary ion mass spectra exclusive to *P. pacificus* and *Ppa-daf-22.1/2* (**d** adults and **g** larvae). Putative chemical assignments (**e** adults and **h** larvae) and structures (**f** adults and **i** larvae) on *P. pacificus* and *Ppa-daf-22.1/2* surfaces (*P*<0.001 by Student's t-test, n=9), present in LIPIDS MAPS (M-H, < 2ppm). **j** Exclusive *P. pacificus* wildtype and *Ppa-daf-22.1/2* mutant chemistries, where phylogeny indicates potential shared regulation of exclusive chemicals and their relative intensity on nematode surfaces.

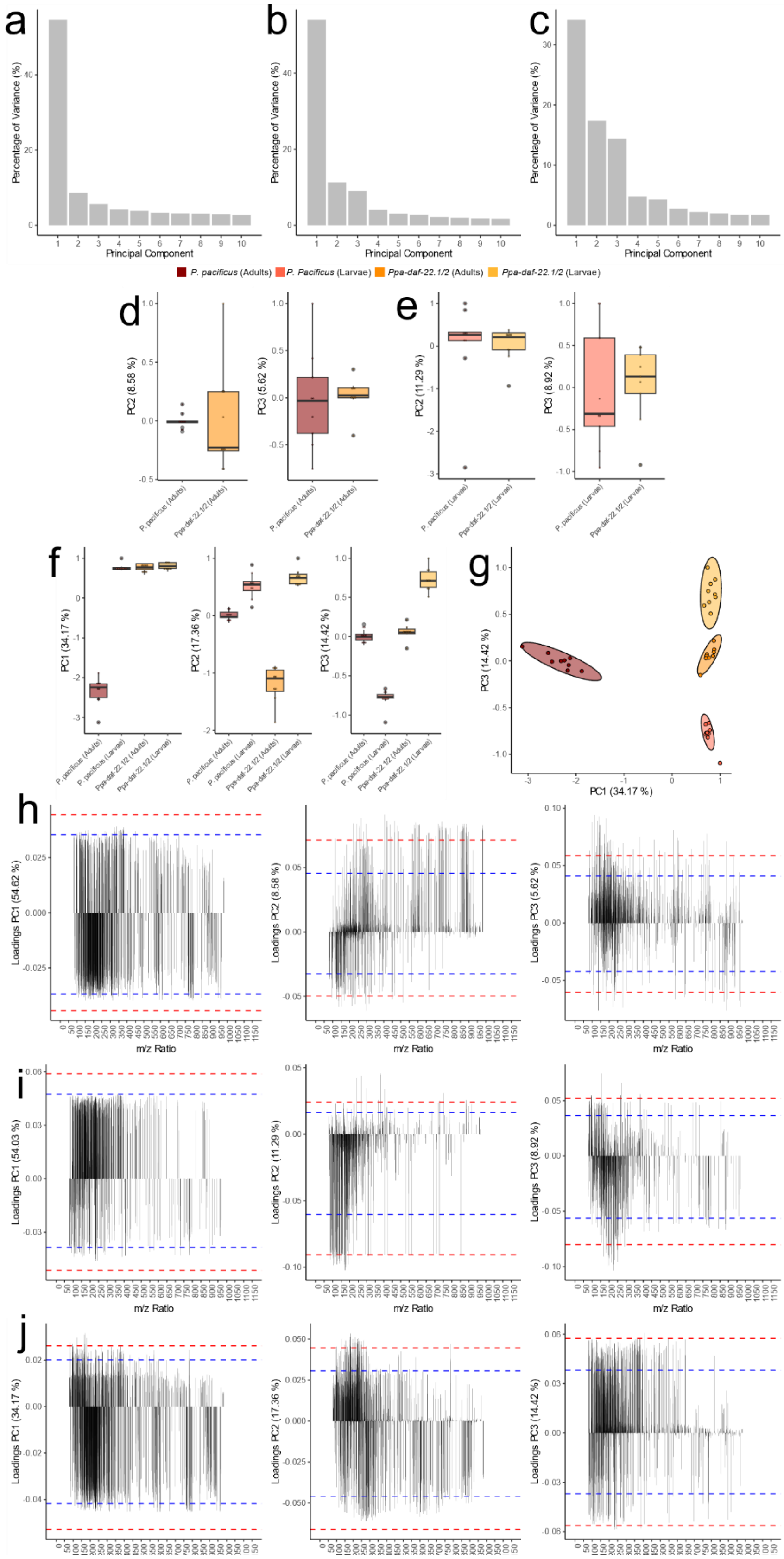


**Fig. S13 | Multivariate analysis of *P. pacificus* wildtype and *daf-22* for each developmental stage.** Percentage of variance on each principal component (**a** adults, **b** larvae and **c** adults & larvae). PC2 and PC3 scores plot (**d** adults and **e** larvae). **f** PC1, PC2 and PC3 wildtype and daf-22 developmental stages scores plot. **g** PC1&3 scores for *P. pacificus daf-22* developmental stages PC1, PC2 and PC3 loadings , where blue and red dashed lines indicate 1 and 2 standard deviations from the mean, respectively (**h** adults, **i** larvae and j adults & larvae)

**Table S1 | Elemental composition ranges of key biomolecules.** A comparative overview of the minimum and maximum counts of key elements in primary biological molecules.

| CATEGORY | GENERAL FORMULA | H | C | N | O | P | S |
|---|---|---|---|---|---|---|---|
| **Carbohydrates** | $Cn(H_2O)n$-1 | 2-12 | 1-7 | 0 | 1-6 | 0 | 0 |
| **Proteins (Amino Acids)** | -(Amino Acid)n | 2-17 | 1-8 | 1 | 1-3 | 0 | 0-1 |
| **Nucleic Acids** | -(Nucleotide)n-1 | 2-9 | 1-8 | 1-3 | 1-6 | 1 | 0 |
| **Fatty Acids & Triglycerides** | $CH_2(CH_2)nCOO$- / $C_{55}H_{97}O_5$ | 4-90 | 2-50 | 0 | 1-4 | 0 | 0 |
| **Phospholipids** | $R_1COOR_2R_3PO_4R_4$ | 8-80 | 4-40 | 1 | 4-8 | 1-2 | 0 |
| **Ceramides** | $C_{34}H_{65}NO_2$ | 5-60 | 9-30 | 1 | 1 | 0 | 0 |
| **Sterols (e.g., Cholesterol)** | $C_{27}H_{45}O$- | 9-44 | 10-25 | 0 | 1 | 0 | 0 |

**Supporting Information Video S1 | *P. pacificus predation*.** Adult *P. pacificus* wildtype predating on *C. elegans* wildtype larvae.

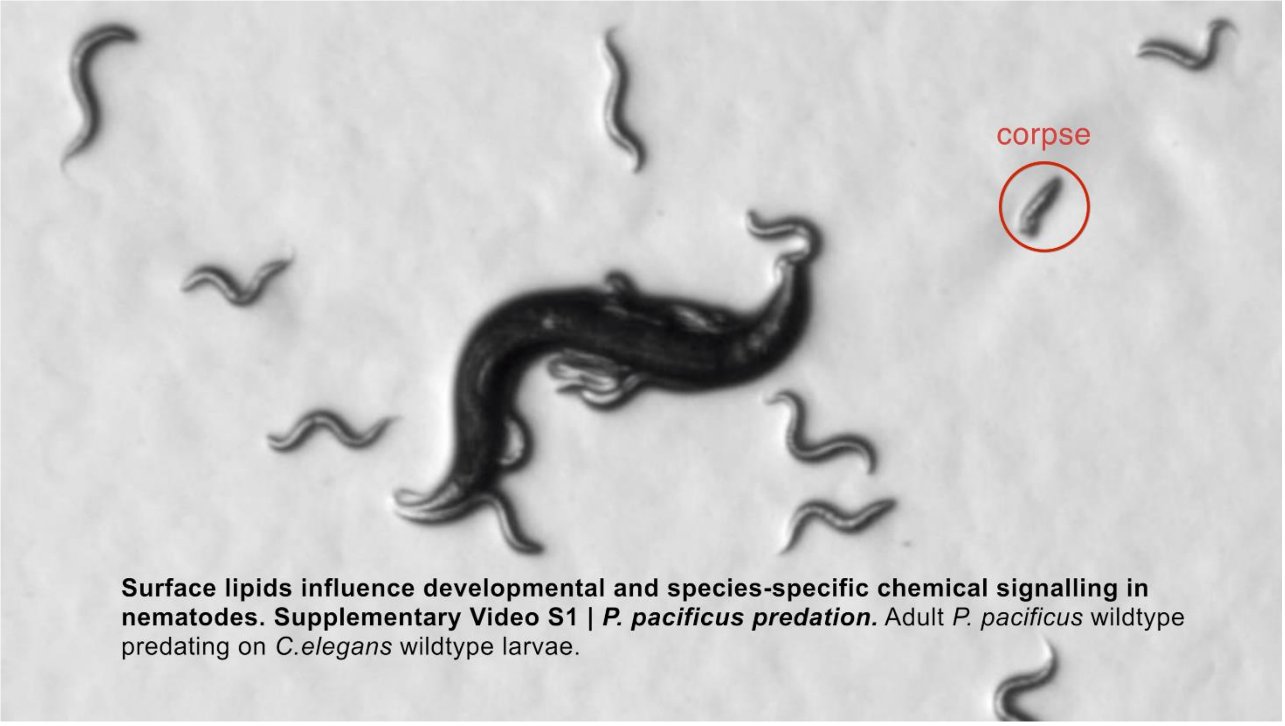
